# RIF1 orchestrates a multi-step restoration of post-replicative H3K9me3

**DOI:** 10.1101/2025.07.14.664777

**Authors:** Naiming Chen, Susanne Bandau, Reshma Ravindran Nair Pushkala Kumari, Bram Prevo, Takayo Sasaki, Shaun Web, Simone di Sanzo, David Shapira, Moritz Voelker-Albert, David M. Gilbert, Constance Alabert, Sara B. C. Buonomo

**Affiliations:** Institute of Cell Biology, School of Biological Sciences, University of Edinburgh, Roger Land Building, Alexander Crum Brown Road, Edinburgh, EH9 3FF, UK; Division of Molecular, Cell & Developmental Biology, School of Life Sciences, University of Dundee, Dundee, DD15EH, UK; San Diego Biomedical Research Institute, San Diego, CA, 92121, USA; Discovery Research Platform for Hidden Cell Biology, University of Edinburgh; MoleQlar Analytics GmbH, Rosenheimer Straße 141h, 81671 Munich, Germany; Peter Gorer Department of immunobiology, King’s College London, London, SE1 9RT, UK

## Abstract

DNA replication causes the dilution of parental histones along with their specific post-translational modifications. The kinetics of restoring these marks on newly incorporated histones dictate how quickly genomic domains regain their epigenetic identity after replication. H3K9me3 is restored extremely slowly; the process of reconstitution, to achieve the pre-replication levels, continues throughout the following G1 phase. The molecular mechanisms behind this slow reconstitution are unknown. We show here that RIF1’s reassociation with heterochromatin during mitotic exit is required to set up a chromatin environment permissive for histone methyltransferases to resume H3K9me3 deposition. RIF1 facilitates the recruitment of SUV39H1, HP1α and HP1β and is required for the increased tri-methylation of H3K9 that occurs during G1 phase. RIF1 is also indispensable for recruiting Protein Phosphatase 1α (PP1α) to heterochromatin, and the interaction between RIF1 and PP1α is essential for the maintenance of H3K9me3 levels. In addition, RIF1-PP1 complex temporally restrains the activity of Aurora kinase at heterochromatin, ensuring that phosphorylation of H3S10 does not precede replication. This creates a time- window permissive for SUV39Hs to initiate the reinstatement of H3K9me3.

## Introduction

Histones are decorated by a combination of post-translational modifications (PTMs) that are closely related to the functional status of the specific genomic segment (Millan-Zambrano et al., 2022). During DNA replication, behind the replication fork, the parental histones, with their specific PTMs, are recycled to daughter strands in a nearly symmetric manner (Petryk et al., 2018, Yu et al., 2018, Gan et al., 2018, Li et al., 2020, Flury et al., 2023). The local re-use of parental histones is the basis for the maintenance of the epigenetic identity, as they work as templates to “copy” the modifications on the naive, newly synthesised and incorporated histones (Scharf et al., 2009, Alabert et al., 2015).

Most epigenetic marks, including DNA methylation, are restored relatively rapidly behind the replication fork, with the newly deposited histones quickly acquiring the same PTMs as the parental histones. H3K27me3 and H3K9me3 are exceptions (Alabert et al., 2015, Reveron-Gomez et al., 2018). When cells enter G1 phase, the extent to which new histones are methylated is still lower than the parental H3K9me3 (Xu et al., 2011) and the recovery of H3K9me3 continues in the following interphase, possibly even in early S-phase (Alabert et al., 2015). Such slow recovery process requires a prolonged period of permissive environment for propagating/depositing these marks, from the moment of DNA replication to the subsequent interphase.

Heterochromatin and euchromatin replication are temporally segregated. In most organisms, heterochromatin replicates during mid-late S phase: the replication timing program controls this temporal separation. It has been proposed that the biological function of the replication timing program is, in fact, to aid epigenetic inheritance (Hiratani and Gilbert, 2009): the sheer delay in firing of the origins of replication embedded within heterochromatin could provide enough time for the Lys- methyltransferases (KMTs) to complete the restoration of H3K9me3 even in early S phase. In support of this hypothesis, it was shown that depletion of the master regulator of replication timing, RIF1 (Cornacchia et al., 2012), during G1 phase induces the loss of H3K9me3 in the following S-phase (Klein et al., 2021).

RIF1 is a large protein that coats heterochromatin, forming large megabase-long domains (RIF1-associated domains, RADs) (Foti et al., 2016). Localised at the nuclear periphery, RIF1 interacts with Lamin B1 (Foti et al., 2016) and is located in the proximity of Lamin A (Roux et al., 2012). A portion of RIF1 is found in the insoluble nuclear fraction (Cornacchia et al., 2012), resistant to nuclease digestion and high-salt extraction, along with the lamina components. In agreement with this, RADs show a high degree of overlap with the Lamina-associated domains (LADs) (Foti et al., 2016). *Rif1* deletion alters the replication timing program in most cell types and organisms (Yamazaki et al., 2012, Seller and O’Farrell, 2018, Peace et al., 2014, Hayano et al., 2012, Cornacchia et al., 2012), with late replicating genomic segments switching to replicating in early S phase and vice versa. In addition, RIF1 is required to limit the interactions between heterochromatin and euchromatin, thereby defining the strength and contributing to the segregation of the A/B compartments (Klein et al., 2021, Gnan et al., 2021). Through its C-terminus, RIF1 interacts with Protein Phosphatase 1a (PP1α) (Sukackaite et al., 2017). This interaction is important for both RIF1-dependent control of chromatin architecture and replication timing (Gnan et al., 2021). PP1 has a key role during mitotic exit: it reverses the phosphorylations by many of the mitotic kinases (CDK, Aurora B and Haspin, reviewed in (Holder et al., 2019)), thus coordinating the resetting of G1 chromatin and the re- assembly of the nuclear envelope (Vagnarelli et al., 2011, de Castro et al., 2017).

H3K9 methylation is catalysed by SET domain containing KMTs. GLP and G9a are responsible for mono and di-methylation of H3K9 (Tachibana et al., 2001, Tachibana et al., 2002, Tachibana et al., 2005), while Suv39H1/2 (Rea et al., 2000) are responsible for depositing H3K9me3. SetDB1 can mono-, di- and tri-methylate H3K9 (Wang et al., 2003). All the H3K9me3 KMTs recognise H3K9me3 (Jurkowska et al., 2017, Wang et al., 2012), SUV39H1/2 via their chromodomains, SetDB1 via its tudor domain, thus enabling the propagation of new tri-methylation to the regions where parental histones are already tri- methylated. In addition, Heterochromatin Protein 1 (HP1), a cofactor that participates to the recruitment and stabilisation H3K9me3 KMTs (Maeda and Tachibana, 2022) binds H3K9me2/3 via its chromodomain (Jacobs and Khorasanizadeh, 2002, Nielsen et al., 2002).

Although the KMTs that generate H3K9m3 in different chromatin contexts are known, the temporal regulation of their function during the cell cycle is still largely unclear. More precisely, the molecular mechanisms dictating the timing of the slow reconstitution of H3K9me3 are unknown.

Aurora B kinase levels oscillate during the cell cycle. Although the peak is reached in G2/M (reviewed in (Willems et al., 2018)), lower but functionally important levels are present in G1 phase (Trakala et al., 2013, Song et al., 2007). Normally, Aurora B phosphorylates H3S10 starting from mid-late S phase throughout G2 (Monier et al., 2007, Hendzel et al., 1997). The presence of H3S10ph interferes with the recognition of H3K9me3 by both HP1 (Fischle et al., 2005, Hirota et al., 2005) and SUV39H1/2 (Rea et al., 2000) chromodomains, promoting their dissociation from heterochromatin. In addition, Aurora B phosphorylates HP1 hinge domain, contributing to the destabilisation of its interaction with chromatin (Terada, 2006). Finally, SUV39H1 is phosphorylated during mitosis by CDK2 and the phosphorylation stimulates its release from heterochromatin (Aagaard et al., 2000, Park et al., 2014). Cell cycle-regulated phosphorylation/dephosphorylation of several of the component of the machinery that maintains H3K9me3 could therefore regulate the timing of H3K9me3 reconstitution after DNA replication. Interestingly, PP1 was identified as a suppressor of variegation (Suv(ar)) gene in *Drosophila melanogaster* (Baksa et al., 1993).

Here we show that the restoration of post-mitotic H3K9me3 levels requires RIF1 first in G1 phase. Eliminating RIF1 in mitotic-arrested cells affects the re-loading of SUV39H1, HP1α, β and of PP1α during mitotic exit, causing the decrease of H3K9me3 levels on newly replicated heterochromatin. We propose that RIF1 re-association with heterochromatin during mitotic exit promotes the assembly of a G1 heterochromatin competent for the H3K9me3 restoration. In addition, we find that the RIF1-PP1 complex is required during S phase to control the timely access of Aurora kinases to heterochromatin. In the absence of a functional RIF1-PP1 complex, H3S10 is phosphorylated ahead of the replication fork, thus potentially further shortening the window of time available for KMTs for completing H3K9me3 in early S-phase and to initiate the restoration of H3K9me3 after replication. In summary, we demonstrate that RIF1 is required for H3K9me3 maintenance throughout the cell cycle, not only by virtue of its role in the control of replication timing: RIF1 plays a role in the reset of the chromatin landscape during mitotic exit, as well as counteracting Aurora kinases at heterochromatin before the passage of the replication fork.

## Results

### RIF1 regulates heterochromatin anchoring to the nuclear periphery

We and others have shown that RIF1 is required for the regulation of chromatin architecture by promoting the partition of the A and B compartments (Gnan et al., 2021, Klein et al., 2021). However, the molecular mechanism through which RIF1 controls chromatin compartmentalisation is unknown (reviewed in (Chen and Buonomo, 2023)). Previous studies (Dan et al., 2014, Klein et al., 2021, Li et al., 2017) have demonstrated that in human cells RIF1 depletion reduces H3K9me3 levels, particularly in RADs transitioning from the B to the A compartment and from late to early replication (Klein et al., 2021). Spracklin and colleagues have also shown that the portion of the heterochromatin that remains unaffected by RIF1 levels is strongly associated with Lamin B1 (Spracklin et al., 2023). Similarly, our findings in mouse embryonic stem cells (ESCs) had shown that not all late replicating heterochromatin is equally dependent upon RIF1. Specifically, RADs highly associated with Lamin B1 maintain their late replication even after *Rif1* deletion, whereas RADs with medium levels of Lamin B1 shift to early replication (Foti et al., 2016). To better understand the features distinguishing RIF1- dependent and independent genomic regions, and to explore how RIF1 influences chromatin organization, we categorised RADs into six clusters, based on their association levels with RIF1 and Lamin B1 across the TAD, used as genomic unit (**Fig. 1A** and **Supp.** Fig.1A). As anticipated, regions in cluster 1 (high RIF1-high Lamin B1) maintain late replication while those in cluster 2 (high RIF1-medium Lamin B1) switch to early replication (**Supp.** Fig. 1B). Interestingly, the compartment association mirrors this pattern: cluster 1 regions maintain the association with the B compartment, whereas cluster 2 regions, already embedded in genomic regions in average with higher eigenvector values (A compartment), loose their B or B-like identity to switch to the A compartment and integrate with their surroundings following *Rif1* deletion (**Fig. 1B** and **Supp.** Fig. 1C). Given that the levels of Lamin B1 association distinguish the regions that are sensitive (cluster 2) and resistant (cluster 1) to *Rif1* deletion, using 3D fluorescent *in situ* hybridisation (FISH) we further investigated, whether RIF1 was differentially required for their association with the nuclear periphery. We find that cluster 2 genomic regions become internalised in Rif1-KO ESCs, while cluster 1 regions maintain the peripheral localisation (**Fig. 1C** and **Supp.** Fig. 1A and 1D). In conclusion, we demonstrate that the regions whose replication timing and association with the B compartment depend on RIF1 are also retained at the nuclear periphery in a RIF1-dependent manner.

**Figure 1.**
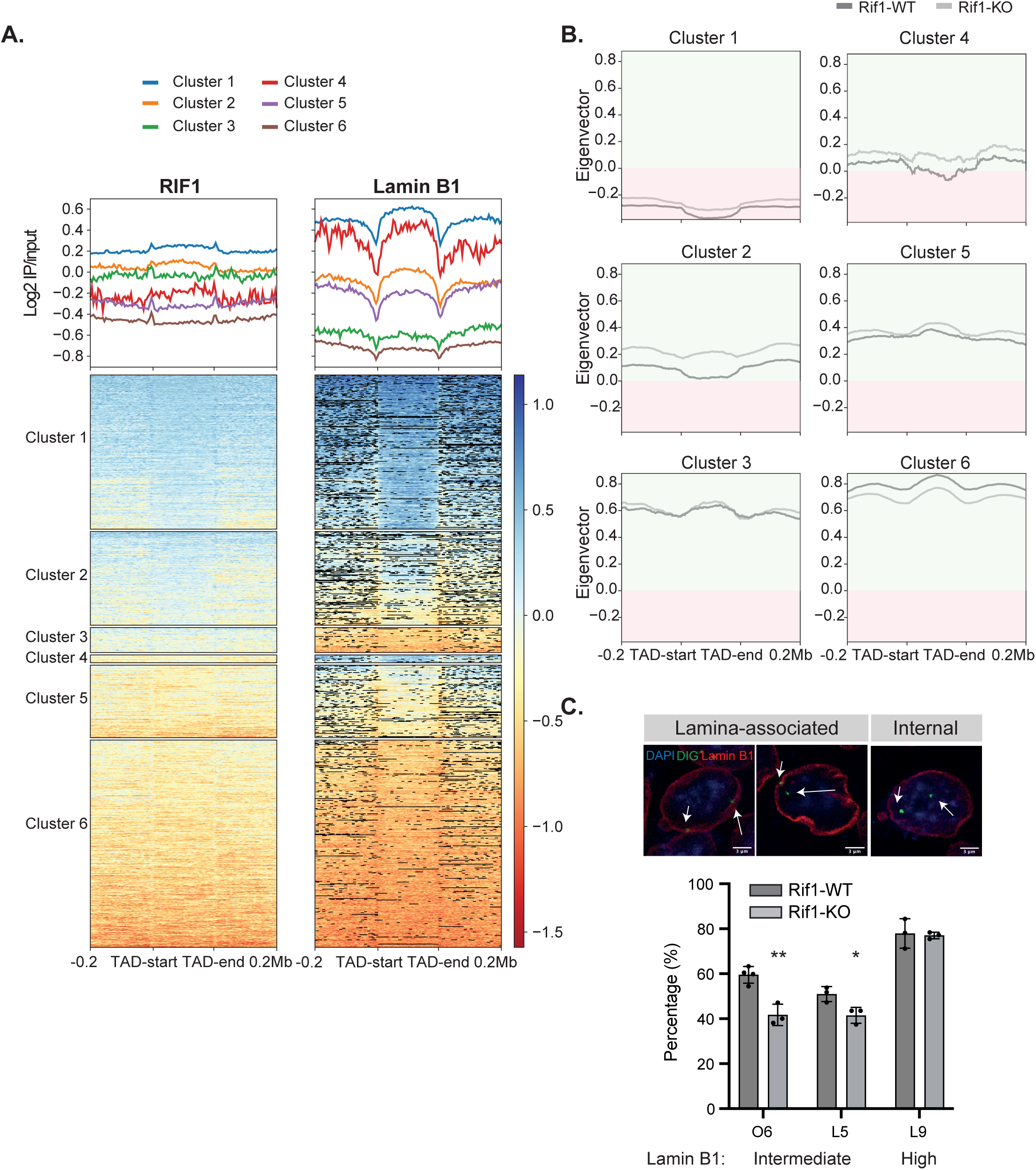
RIF1 is essential for the peripheral localisation and association with the B compartment of heterochromatin bound by intermediate levels of Lamin B1. A. Heatmaps showing TADs clustered based on RIF1 (ChIP-seq) (Foti et al., 2016) and LaminB1 (Dam ID, Peric-Hupkes et al., 2010) levels. Flanking regions of ±0.2 Mb were included around the start and end of each TAD. The ChIP and DamID signals were averaged over 1kb windows. The TADs were calculated from HiC data from (Gnan et al., 2021). Clusters 1, 2, and 3 are RIF1 high, with high, intermediate, low level of Lamin B1 respectively. Clusters 4, 5, 6 are RIF1 low, with high, intermediate, and low Lamin B1 respectively. The 6 clusters were obtained by k-means clustering. B. A and B compartment association analysed in the 6 clusters defined in A. The eigenvector values were averaged over 1kb genomic windows. Rif1-WT=dark grey; Rif1-KO=light grey. Rif1- KO cells are *Rif1^flox/flox^ Rosa26^CreERT/+^* cells treated for 4 days with 4-hydroxytamoxifen (OHT) to activate the CRE recombinase and delete *Rif1*. Rif1-WT are *Rif1^+/+^ Rosa26^CreERT/+^* cells treated for 4 days with 4-hydroxytamoxifen (OHT) (see Materials and Methods). The A compartment is shadowed in green, the B in red. C. Representative images of Lamin B1 immunofluorescence combined with 3D-FISH (IF-3D-FISH) of selected genomic regions. DAPI(Blue), Lamin B1 (Red) and FISH probe (green) are shown. The FISH signal is highlighted by the arrows. For each probe, nuclei were assigned to the category of “lamina-associated” when at least one allele was located to the nuclear periphery (see Material and Methods), while nuclei where both alleles were not at the nuclear periphery were counted in the category of “internal localisation”. The IF-3D-FISH was quantified to determine the peripheral association of the indicated regions. The percentage of nuclei showing at least one lamina-associated signal was plotted. Each point represents an independent Rif1-WT or Rif1-KO clone. At least 80 cells per sample were counted. The mean and standard deviation are shown in the plot. The statistical analysis was performed with Welch’s *t*-test, p<0.05*. p<0.01**, p<0.001***.

### Upon RIF1 loss, the decrease of H3K9me3 is not the main driver of chromatin architecture or replication timing changes

Do the same affected genomic regions depend on RIF1 to regulate H3K9me3 levels? H3K9me2/3 is crucial in regulating chromatin architecture (Feng et al., 2020) and, specifically, in modulating the anchoring of heterochromatin to the nuclear lamina (Towbin et al., 2012, Armstrong et al., 2020, Poleshko et al., 2019). In mouse ESCs lacking both KMTs *Suv39h1* and *Suv39h2* (Suvdn)(Lehnertz et al., 2003), H3K9me3 levels are reduced within cluster 2 genomic regions (**Supp.** Fig. 2A). These same regions also shift towards the A compartment and become internalised, similarly to what we have observed in Rif1-KO cells (**Supp.** Fig. 2B, C **and D**). This suggests a potential link between *Rif1* deletion, compartment shifting from B to A and the internalisation of cluster 2 genomic regions via diminished H3K9me3. To test this hypothesis, we analysed H3K9me3 levels and distribution by ChIP-seq in Rif1-KO ESCs. We found that *Rif1* deficiency affects H3K9me3 levels across both cluster 1 and 2 RADs (**Fig. 2A and Supp.** Fig. 3A and B), although only cluster 2 regions undergo the B-to-A compartment shift **(Fig. 1B)** and internalisation (**Fig.1C and Supp.** Fig. 1A and D).

**Figure 2:**
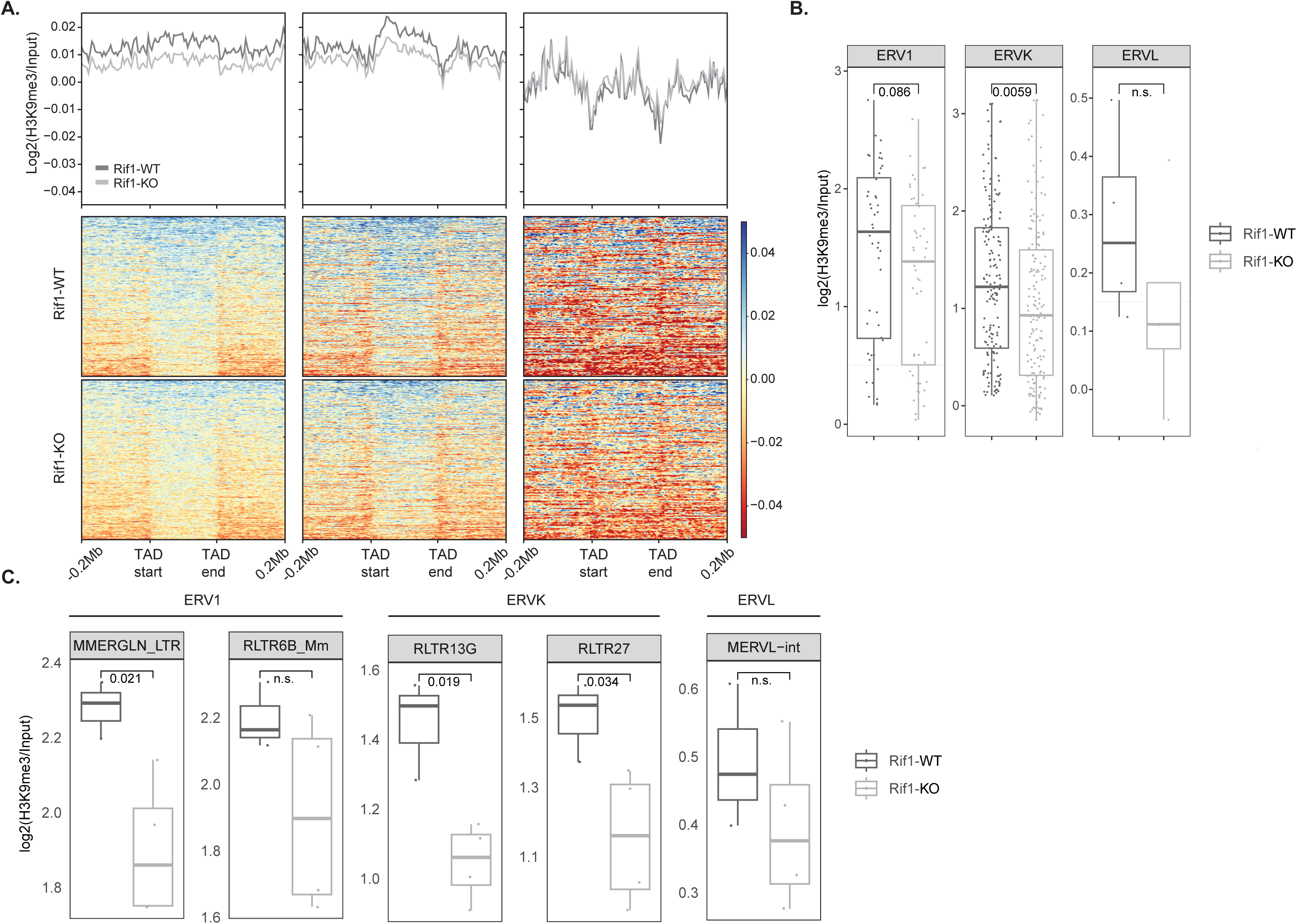
Rif1 deficiency decreases H3K9me3 across RADs. **A.** Comparison of H3K9me3 ChIP signals averaged over 5kb windows, across three RIF1-high clusters (clusters 1, 2, and 3) between Rif1-WT and Rif1-KO ESCs. **B.** ChIP-seq quantification of H3K9me3 levels across the three ERV classes, pre-selecting ERVs with a log2(H3K9me3_WT/Input) >0.1. ERVK= ERV2; ERVL=ERV3. In the cell lines and conditions used in this study (see Materials and Methods), we find that ERVLs are only modestly enriched for H3K9me3 and are consequently under-represented in this analysis. The *p-*values have been calculated by Wilcoxon rank sum test. **C.** ERVKs (RLTR13G and RLTR27) exhibit the most significant decrease in H3K9me3 levels following *Rif1* deletion. A reduction in H3K9me3 is also observed in ERVL (MERVL-int) and ERV1 (MMERGLN_LTR and RTR6B_Mm) elements, although the differences between Rif1-WT and Rif1-KO do not always achieve statistical significance. *p*-values were calculated using a *t*-test.

Consistent with previously published data (Li et al., 2017), we find that H3K9me3 is particularly reduced at endogenous retroviruses (ERVs) (**Fig. 2B**), especially class 2 ERVs, ERVKs (**Fig. 2C**). ERVKs are enriched in cluster 1 regions (**Supp.** Fig. 3C) and show the highest enrichment for RIF1 among ERVs (**Supp.** Fig. 3D), although other repeat classes also have significant RIF1 enrichment (**Supp.** Fig. 3E). The loss of H3K9me3 peaks in Rif1- KO cells corresponds with RIF1 peaks across most clusters (**Supp.** Fig. 3F), suggesting a direct role of RIF1 in the maintenance of H3K9me3.

These data also show that while in cluster 2 regions H3K9me3 levels, replication timing and nuclear architecture all change upon loss of RIF1, in cluster 1 regions *Rif1* deficiency induces a loss of H3K9me3 comparable to the loss in cluster 2, but this is not accompanied by release of the anchoring from the lamina, loss of association with the B compartment or switch to early replication timing. The regulation of H3K9me3 by RIF1 is therefore independent of its control over chromatin architecture and replication timing, at least in cluster 1 regions.

In a complementary experiment, we found that artificially restoring peripheral localisation and H3K9me3 levels without RIF1 cannot reinstate late replication timing. We established an inducible system to re-tether one of the cluster 2 region, O6, back to the lamina in Rif1-KO cells (**Supp.** Fig. 4A). In brief, we specifically targeted multiple copies of a Tetracyclin operator (*TetO*) to one of the two alleles of O6 (**Supp.** Fig. 4B, see Materials and Methods for details). In the same cells, we expressed a mini-*Lap2β* ((Foisner and Gerace, 1993, Finlan et al., 2008), a Lamin B1-interacting protein) fused to *Frb* and a Tetracyclin Repressor (*TetR*) fused to *Fkbp* (**Supp.** Fig. 4C). FRB-miniLAP2β was efficiently associated with the nuclear lamina (**Supp.** Fig. 4D**, top**) and binding of the TetR-FKBP to the *TetO* did not alter the late replication timing of the region (**Supp.** Fig. 4H**, WT_DMSO**). Upon Rapamycin treatment, FKBP and FRB dimerise, pulling the TetO-containing O6 region to the nuclear lamina (**Supp.** Fig. 4D**, middle and bottom**). Tethering back to the nuclear periphery (**Supp.** Fig. 4E and F) and restoration of the H3K9me3 levels **(Supp.** Fig. 4G) are not sufficient to restore late replication and TetO-O6 remains early replicating in the absence of RIF1(**Supp.** Fig. 4H, KO_Rapa).

### RIF1 promotes the restoration of H3K9me3 levels in G1 phase

It has been proposed that the changes of replication timing caused by RIF1 deficiency could be indirectly responsible for the decrease of H3K9me3 (Klein et al., 2021). However, our observation that H3K9me3 decreases also in cluster 1 regions that remain late replicating suggests that factors other than replication timing must additionally contribute to H3K9me3 maintenance. What is the molecular mechanism by which RIF1 controls H3K9me3? H3K9me3 restoration continues from one cell cycle into the next, throughout the next G1 phase (Alabert et al., 2015). RIF1 progressively reassociates to anaphase chromosomes, coincidentally with the dephosphorylation of H3S10ph (**Supp.** Fig. 5A), which is a likely prerequisite for resuming H3K9me3 restoration (Rea et al., 2000, Peng et al., 2018, Hirota et al., 2005, Fischle et al., 2005). This timing suggests that the reassociation of RIF1 with heterochromatin during mitotic exit could be important for restarting the process of restoration of H3K9me3. To test this hypothesis, we generated an ESC line where both *Rif1* alleles were tagged with miniAID-mCherry (**Supp.** Fig. 5B), allowing 5-ph-IAA-induced degradation (Saito and Kanemaki, 2021) (**Supp.** Fig. 5C and D). With this tool, we could induce the acute depletion of RIF1 in cells arrested in mitosis (when RIF1 is least chromatin bound), to then released the cells into the following G1 phase (**Fig. 3A** and **Supp.** Fig. 6A) in the absence of RIF1 (**Supp.** Fig. 6B and C). In these experimental conditions, cells undergo S phase in presence of RIF1 and of a functional replication timing (**Fig. 3B**). Consequently, any eventual effect of RIF1 deficiency on H3K9me3 levels can only be attributed to a metaphase-to-G1-specific RIF1 function. RIF1 degradation did not affect the release from the arrest and the re-entry into G1 phase observed by FACS (**Supp.** Fig. 6A). By iPOND-SILAC mass spectrometry we found that on both parental and newly incorporated histones H3K9me3 levels were mildly reduced in the ESCs where we had induced RIF1 degradation for 4 hours (**Fig. 3C and D** and **Supp.** Fig. 6D). Importantly, the lower H3K9me3 levels were also accompanied by a tendency to accumulate H3K9me2 (**Fig. 3C, D and Supp.** Fig. 6E). Considering that the ChIP-seq analysis shows that RIF1 is required for H3K9me3 maintenance only in a specific subset of the genome and that not all the cells are released from the G2/M arrest (**Supp.** Fig. 6A), the limited extent of the effect was expected. Interestingly, the reduced levels of H3K9me3 are specific to the peptides carrying the K9me3 individual modification. The peptides where, in addition, K14 is acetylated do not show the same tendency (**Supp.** Fig. 6F, G **and H**). In mouse embryonic fibroblasts, these peptides are specific to histones H3 that occupy the Long Interspersed Nuclear Elements (LINEs) and are modified by a subset of SetDB1 (Jurkowska et al., 2017). Our analysis also showed that H3K37me3 and H3K36me3 were not affected (**Supp.** Fig.6I **and J**), suggesting that RIF1 is not involved in the restoration of these marks. In summary, we propose that RIF1 is required during mitotic exit to reinitiate the restoration of H3K9me3 levels in G1 phase, suggesting a role for RIF1 that transcends its functions during S phase.

**Figure 3.**
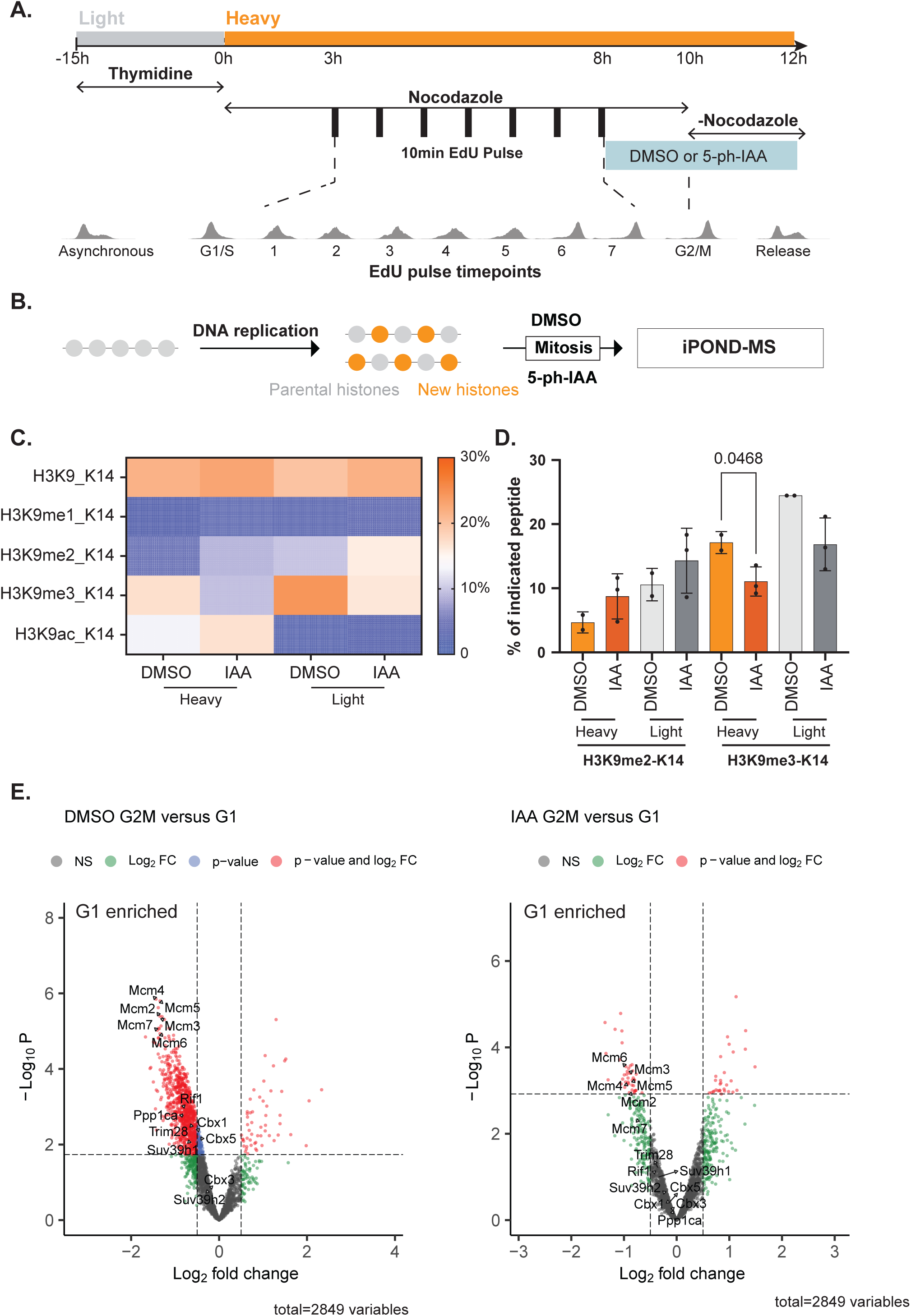
Degradation of RIF1 during mitosis impairs H3K9me3 restoration in the following G1. **A.** Experimental design: cells grown in medium supplemented with a light- isotope-labelled Arginine/Lysine (R0K0) were synchronised at the G1/S boundary by thymidine treatment for 15 hrs, then released for 8 hrs in S phase in media containing heavy isotope-labelled Arginine/Lysine (R10K8) and nocodazole. Mid-late replicating chromatin was pulse-chase labelled for 10 min. with EdU at the indicated timepoints. After 8 hrs from the release, 5-ph-IAA was added for 2 hrs to degrade RIF1. Cells were released from nocodazole into the next G1 for additional 2 hrs in presence of 5-ph-IAA. Pulse-labelled cells were then collected and pooled for iPOND-SILAC mass spectrometry. See also Additional information Figure 1. **B.** Schematic of the outcome of the experiment outlined in A. Old and newly synthesized histones are labelled with light R0K0 (grey) and heavy R10K8 (orange) isotopes respectively and incorporate during a normal S-phase, that takes place in presence of RIF1. RIF1 degradation is induced only during mitosis. Mid-late replicating chromatin in cells that underwent mitosis and entered G1 in the presence (DMSO) or absence (5-ph-IAA) of RIF1 was isolated by iPOND and histones were extracted for mass spectrometry (MS) analysis. **C.** Heatmap of the average percentage (DMSO n=2; IAA n=3) of indicated modified peptides (H3K9me0/1/2/3/ac_K14) over the total H3 peptide aa 9-17. Both parental (light) or newly incorporated (heavy) histone H3 extracted from cells in G1 were analysed. **D.** The barplot shows the percentage of H3K9me2_K14 and H3K9me3_K14 relative to the total histone H3 peptide aa 9-17. DMSO (n=2), IAA (n=3). The mean and standard deviation are shown. *p*-values were calculated using a Welch’s *t*-test. **E.** Experiment design similar to A, but no heavy R/K was used. See also Additional information Figure 2. Volcano plots summarising the efficiency of G1 re-association of proteins to mid-late replicating chromatin, relative to G2/M. Data show iPOND-MS analysis of cells treated with DMSO (left) or 5-ph-IAA (right) from four independent experiments. The log_2_ fold change indicates the difference between G2/M and G1. The horizontal cutoff shows the adjusted *p*-value at 0.05 and vertical cutoffs indicate the log_2_ fold change at ± 0.5.

### RIF1-PP1 is required for H3K9me3 restoration

Li and colleagues have shown that RIF1 is necessary for the recruitment of SUV39H1 to various ERVs (Li et al., 2017). In agreement with this, we find that the selective removal of RIF1 during mitosis affects the normal heterochromatin reassociation in G1 not only of SUV39H1, but also of HP1α and β, two H3K9me3 readers and recruiters of SUV39H1 (**Fig. 3E** and **Supp.** Fig. 7A). Even more strikingly, we discovered that RIF1 depletion causes a severe reduction of the re-association of PP1α to G1 heterochromatin, pointing to RIF1 being a central recruiter of PP1α to mid-late replicating heterochromatin (**Fig. 3E and Supp.** Fig. 7A). Finally, the re-loading of the MCM complex onto the G1 chromatin is also slightly reduced. This observation is in agreement with RIF1-PP1 requirement to reverse the mitotic phosphorylation of ORC1 and for efficient origin licencing (Hiraga et al., 2017). To investigate if RIF1-PP1 interaction is important for the restoration of H3K9me3 levels, we analysed cells where the sole source of RIF1 is a mutant knock-in allele, carrying two point mutations that abolish the interaction with PP1α (Gnan et al., 2021). In these cells we found that, similar to the effect of *Rif1* deletion compared to wild type cells (**Fig. 2A and B and Supp.** Fig. 3A), the loss of RIF1-PP1 interaction reduces the levels of H3K9me3 at various ERVs compared to the matched hemizygote control (Rif1-HEMI, **Fig. 4A, B and Supp.** Fig. 8A). Similarly, we found that loss of RIF1-PP1 interaction also induces a decrease of H3K9me3 levels in cluster 1 and 2 genomic regions (**Fig. 4C and Supp.** Fig. 8B) and, more specifically, within the O6 and L5 regions, that are internalised in the Rif1- KO ESCs (**Supp.** Fig. 8C). However, these regions are internalised at a similar frequency in Rif1-HEMI and Rif1-ΔPP1 cells (**Supp.** Fig. 8D), in addition to displaying a similar shift to the A compartment (**Supp.** Fig. 8E). Since in both Rif1-HEMI and Rif1-ΔPP1 RIF1 is present at half dosage compared to the wild type (Gnan et al., 2021), this result suggests that RIF1 dosage, rather than the decrease of H3K9me3, is the driving force behind the remodelling of nuclear architecture in the Lamin B1 intermediate regions (cluster 2).

**Figure 4:**
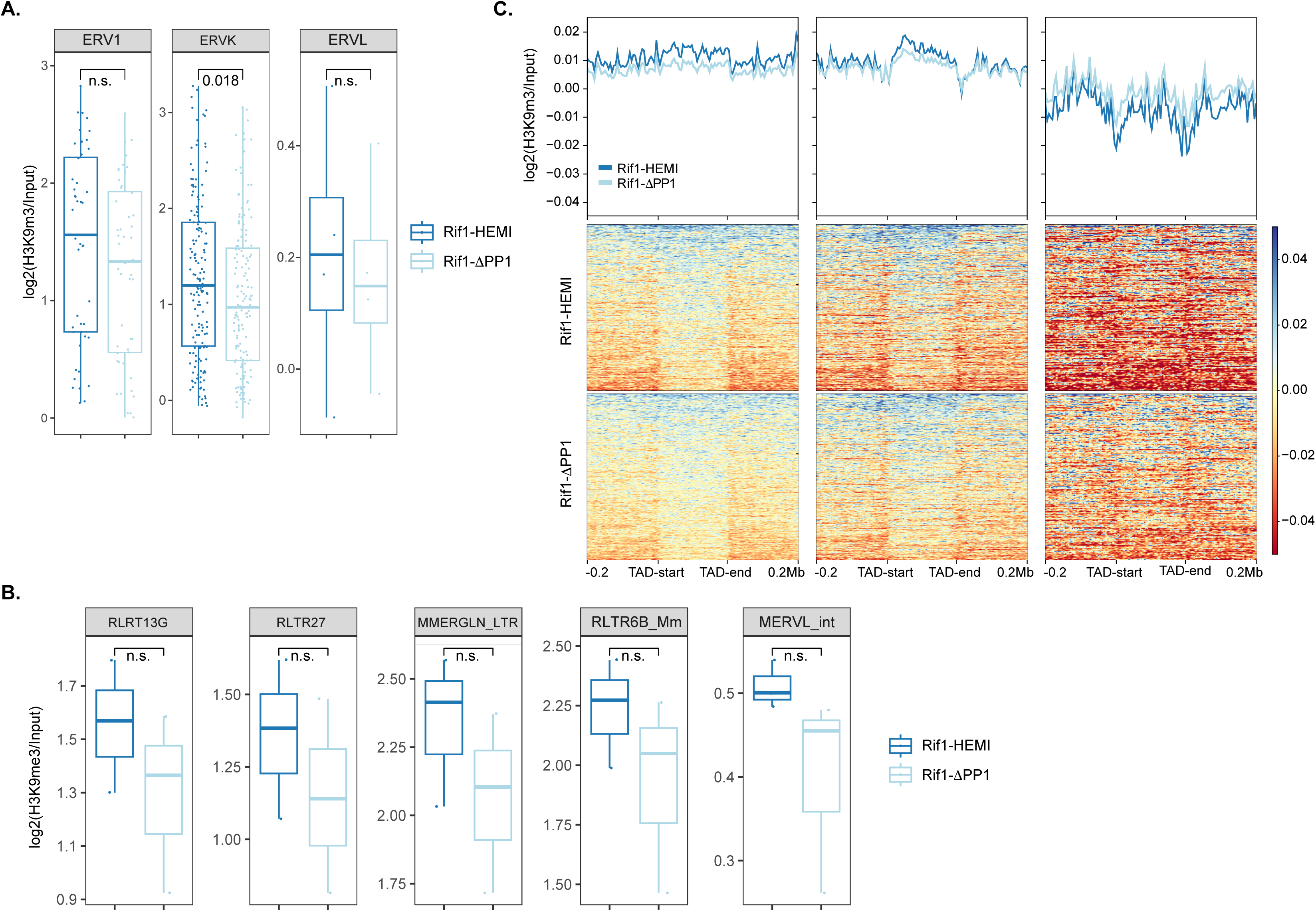
Loss of RIF1-PP1 interaction leads to H3K9me3 reduction across RADs. **A.** Chip-seq quantification of H3K9me3 levels across ERVs with a log2(WT/Input) >0.1 (as in Figure 2) in Rif1-HEMI and Rif1-ΔPP1. The *p-*values have been calculated by Wilcoxon rank sum test. Treatment with OHT of the *Rif1^flox^*^/ΔPP1^ *Rosa26^CreERT/+^* cell lines induces the deletion of the functionally wild type copy of *Rif1 (Rif1^flox^* allele), leaving the second allele, Rif1-ΔPP1 minigene, as the only, hemizygous source of RIF1 (see Materials and Methos and Gnan et al., 2021). The control for these cells is the Rif1-HEMI cell line (*Rif1^flox^*^/TgWT^ *Rosa26^CreERT/+^*), build in the same way, except that the mini-gene encodes for wild type RIF1 (targeted wild type=TgWT). **B.** Changes in H3K9me3 levels (expressed as log2(H3K9me3/Input)) across representative ERVs from all three classes. Similar to *Rif1*- KO cells shown in Fig. 2, cells with impaired RIF1-PP1 interaction (Rif1-ΔPP1) exhibit a tendency to reduce H3K9me3 levels over these ERVs that are occupied by RIF1 (Supp. Fig. 3C). **C.** Comparative analysis of H3K9me3 ChIP signals averaged over 5kb windows in the 3 RIF1-high clusters (clusters 1,2 and 3). Rif1-HEMI and Rif1-ΔPP1 are shown. *p*-values were calculated using a *t*-test. A general higher variability among the Rif1-HEMI clones reduces the power of statistical analysis in these samples. Nevertheless, Rif1-ΔPP1 cells display the tendency to lower levels of H3K9me3 compared to the control Rif1-HEMI.

In conclusion, we have shown that RIF1 is required for the heterochromatin reassociation in G1 of SUV39H1, HP1α, HP1β and PP1α, all of which are heterochromatin proteins required for the restoration of H3K9me3 levels. We also found that RIF1-PP1α interaction is required for the maintenance of H3K9me3 levels. As both SUV39H1 (Aagaard et al., 2000, Park et al., 2014) and HP1α (Terada, 2006) are phosphorylated during mitosis to be released from chromatin, we hypothesise that one or both could be dephosphorylated by RIF1-PP1α during mitotic exit to promote their reassociation with heterochromatin.

### RIF1-PP1α time the activity of Aurora kinases on heterochromatin

In the course of this study, we observed an increased percentage of cells with H3S10ph foci in Rif1-KO and Rif1-ΔPP1 ESCs (data not shown). To determine whether this increase was due to the previously reported accumulation in the G2 phase (Gnan et al., 2021) or a change in Aurora kinases activity, we pulsed Rif1-WT, Rif1-KO, Rif1-HEMI and Rif1-ΔPP1 ESCs with EdU. We then used FACS to sort G1 (**Fig. 5A**, P1) and early S phase (**Fig. 5A**, P2) populations, based on the DNA content. Subsequent fixation and staining reveal a slightly higher percentage of cells with H3S10ph foci already present in G1 phase in both Rif1-KO and Rif1-ΔPP1 cells (**Fig. 5B**, P1 and **C**, top). This is in agreement with the observation that after inducing RIF1 degradation, we still find H3S10ph positive cells in telophase, suggesting a delay in the removal of a portion of H3S10ph (**Supp.** Fig. 5C, arrows). The percentage of positive H3S10ph cells further increases in early S phase (**Fig. 5B**, P2 and **C**, bottom), well before the phosphorylation of S10 on histone H3 by Aurora B normally starts becoming detectable.

**Figure 5.**
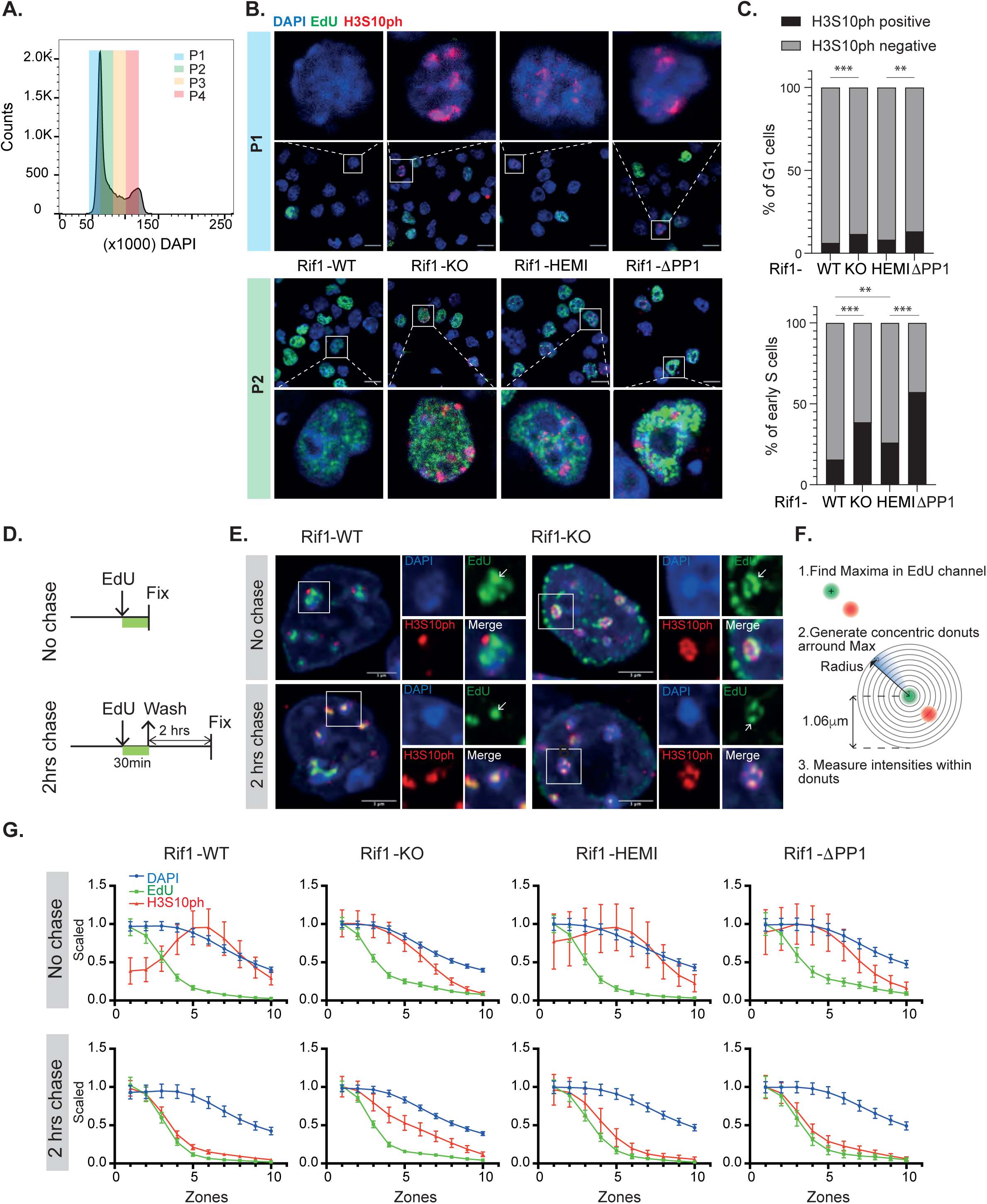
***Rif1* deficiency leads to aberrant dynamics of H3S10ph in both G1 and S phase. A.** Cells pulsed for 30 min. with EdU were fixed and sorted, based on the DNA content, in P1 (G1), P2 (early S), P3 (mid S) and P4 (late S) fractions. Example of the sorting profile**. B.** Representative images of immunostaining for H3S10ph combined with EdU staining on sorted cells from P1 and P2 fractions. DNA=DAPI, blue; EdU=green; H3S10ph=red. **C.** Quantification of H3S10ph foci in P1 and P2 fractions from three independent experiments. In P1, G1 nuclei were identified as EdU negative. In P1 and P2, early S cells were identified by a diffuse EdU signal. The plot shows the of percentage of nuclei with H3S10ph foci. Sample size of P1: WT=572; KO=805; HEMI=815; ΔPP1=839. Sample size of P2: WT=436; KO=256; HEMI=387; ΔPP1=187. Statistical analysis was performed by Fisher’s exact test. **D.** Experimental design: cycling cells of different genotypes were either pulsed with EdU and fixed or pulsed with EdU and chased for 2 hrs before fixation, followed by immunostaining for H3S10ph combined with EdU staining. **E.** Representative images from Rif1-WT and KO cells. Chromocenters are an easily identifiable nuclear domain where the order: replication-followed-by-H3S10 phosphorylation is well characterised. We have therefore focused our analysis on/around chromocenters. As expected, in Rif1-WT, EdU (green) and H3S10ph (red) do not co-localise (No Chase). However, 2 hrs after the EdU pulse, H3S10ph is observed on the chromocenter that was replicated 2 hrs prior (2 hrs Chase), confirming that the phosphorylation of H3S10 normally follows the passage of the replication fork. DNA=DAPI, blue. **F.** Illustration of the image analysis pipeline to measure H3S10ph signals relative to EdU foci. The most prominent EdU foci/spots were identified. Then H3S10ph signals were measured within ten concentric circles centred on EdU foci. **G.** Quantification of H3S10ph position with respect to EdU foci, as described in F. Three clones of each genotype were used for the quantification. The plots show the mean and 95% confidence intervals of DAPI, EdU and H3S10ph. An independent experiment is shown in Additional information Fig.3.

Under normal conditions, Aurora B binds to heterochromatin on the wake of the replication fork (Alvarez et al., 2023). The RIF1-PP1α complex loses the heterochromatin enrichment prior to the arrival of the replication fork (Seller and O’Farrell, 2018, Cho et al., 2022). RIF1-PP1α could therefore shield heterochromatin substrates from early Aurora kinase activity, until replication is underway. In case of impaired recruitment of PP1α at heterochromatin, such as in Rif1-KO or Rif1-ΔPP1 cells, H3S10 could become a premature substrate for Aurora kinase. Supporting this hypothesis, both the slight G1 and the more substantial early S phase increase of H3S10ph in Rif1-KO cells could be reversed by treatment with an Aurora kinases inhibitor, hesperadin (**Supp.** Fig. 9A). We found no evidence that RIF1 deficiency affects the levels or the dynamics of Aurora B accumulation during the cell cycle (**Supp.** Fig. 9B and C)

The precocious appearance of H3S10ph in RIF1-deficient cells does not necessarily imply that the order of the events (replication first, H3S10ph after) is altered. H3S10ph could still be deposited post-replication in regions of heterochromatin abnormally replicated early in S phase, due to *Rif1* deficiency. We set out to test if lack of functional RIF1 subverts the order of the events, causing the replication fork to go through heterochromatin where H3S10 has been precociously phosphorylated. To this end, we performed a pulse-chase experiment (**Fig. 5D**). Typically, in wild type cells, H3S10ph only accumulates at pericentromeric heterochromatin post-replication (**Fig. 5E, F, G and Additional information Fig. 3**). However, in Rif1-KO and Rif1-ΔPP1 mESCs, replication forks and H3S10ph overlap at/around chromocenters (**Fig. 5E, F, G and Additional information Fig. 3**). Moreover, we found that even yet-to-be replicated pericentromeric heterochromatin, identified as MCM3-associated EdU negative chromocenters (Yan et al., 1993, Aparicio et al., 2012), harbours H3S10ph in the absence of functional RIF1 (**Supp.** Fig. 9D and E). These findings underscore the essential role of RIF1-PP1α in binding to heterochromatin: it is crucial not only for delaying the firing of replication origins, perhaps allowing adequate time for H3K9me3 reconstitution prior to replication, but also for counteracting the premature activity of Aurora B that interferes with H3K9 tri- methylation by phosphorylation of H3S10.

## Discussion

The discovery of heterochromatin’s uniquely slow post-replicative reconstitution dynamics (H3K9me3 and H3K27me3) has been puzzling. Intuitively, “weakened” heterochromatin might give rise to unwanted gene expression. Perhaps, delaying heterochromatin replication until the end of S phase could act as a safeguard: by the time post-replicative chromatin reacquires full transcriptional competence (Stewart-Morgan et al., 2019), the chromosome condensation occurring in preparation for mitosis and reduced chromatin accessibility could lower the risks. However, the biological reasons for the evolution of such slow dynamics in H3K9me3 and H3K27me3 reconstitution are still unclear. One theory suggests that this delay creates a window of opportunity for cell fate changes in G1 phase.

From a molecular perspective, the mechanisms underlying this delay are also unclear: Klein and colleagues have shown that eliminating RIF1 in late G1 phase changes replication timing before H3K9me3 levels drop within the same cell cycle, leading to the hypothesis that anticipated firing of origins of replication within heterochromatin is responsible for the defective H3K9me3 reconstitution. This suggests the hypothesis that the biological function of the replication-timing program, a longstanding mystery, might be to help the transmission of the epigenetic information across S phase.

In this study, we unveil a complex role for RIF1 in promoting H3K9me3 transmission, that transcends the sole regulation of replication timing.

First, RIF1 binding to heterochromatin during mitotic exit aids in the reconstitution of a G1 chromatin permissive for H3K9me3 deposition, by promoting the re-association of HP1α, HP1β and SUV39H1. Our mass spectrometry analysis of the G1 proteome reveals that RIF1’s failure to reassociate with chromatin during mitotic exit causes a generalised delay in the re-assembly of G1 heterochromatin. Furthermore, RIF1 serves a key recruiter of PP1α at mid-late replicating chromatin. The delayed/inefficient re-assembly of the molecular machinery responsible for reinstating H3K9me3 could be part of a more general requirement for RIF1-PP1α to reverse mitotic phosphorylation and re-build G1 chromatin. The less efficient reconstitution of H3K9me3 likely results from delayed dephosphorylation of a number of proteins. Although testing this hypothesis is technically challenging due to the lack of reliable antibodies specifically recognizing the phosphorylated proteins, and even though dephosphorylation is probably delayed rather than completely halted, our results indicate that RIF1 deficiency leads to the delayed removal/premature appearance of H3S10ph during telophase and G1 phase. Importantly, as only a subset of RADs is strictly dependent upon RIF1, during mitotic exit RIF1 is likely to be absolutely required only for the dephosphorylation of the PP1-target proteins associated with these RADs. Other RIPPOs (Regulatory Interactors of Protein Phosphatase One) are associated with heterochromatin, probably in different domains non-overlapping or only partially overlapping with RIF1-dependent RADs, performing a similar function. For example, Repo-Man/PP1 (β or γ), associated with the nuclear periphery, has a key role in dephosphorylating H3S10ph during mitotic exit (Vagnarelli et al., 2011) and is instrumental for the re-assembly of G1 chromatin and for maintaining H3K9me3 (de Castro et al., 2017). KI-67, another RIPPO for PP1β and γ, localises at LADs (van Schaik et al., 2022) and participates to the maintenance of H3K9me3 (Sobecki et al., 2016).

Next, we show that the presence of the RIF1-PP1α complex on heterochromatin limits Aurora kinase activity to a time window that follows the replication fork’s passage. Normally, the association of Aurora B with newly replicated heterochromatin and the consequent phosphorylation of H3S10 initiates chromatin condensation, probably pausing H3K9me3 reconstitution.

By delaying the phosphorylation of Ser10 on histone H3, and, perhaps, by simultaneously delaying the firing of heterochromatin’ origins of replication, RIF1-PP1 could provide SUV39H1 with sufficient time to complete H3K9me3 deposition prior to the passage of the replication fork. If H3S10ph appears on heterochromatin prior to replication (in G1 or early S phase), the completion of K9 tri-methylation on both parental and newly incorporated H3 prior to the new round of replication would be impaired. Moreover, there would be no time window available for SUV39Hs to initiate methylation of the newly incorporated histones post-replication.

Our data suggest that the molecular mechanism of the clock behind the unique dynamics of the reconstitution of H3K9me3, is a multistep process involving PP1 to both reverse and counteract kinases’ activity to re-assemble heterochromatin and maintain its identity.

Regarding the implications of reduced H3K9me3 maintenance, we had hypothesised that the changes of chromatin organisation in the absence of functional RIF1 could be linked to RIF1’s role in the maintenance of H3K9me3. In this work, however, we demonstrate that the decrease of H3K9me3 caused by *Rif1* deletion cannot account alone for the shifts from the B towards the A compartment or for the internalisation of heterochromatin. Chromatin architecture in cluster 2 regions is exquisitely sensitive to RIF1 dosage. We reveal that the full complement of RIF1 is specifically required for anchoring these regions to the nuclear periphery, suggesting that the changes in chromatin three-dimensional organisation that follow the partial or full loss of RIF1 could perhaps be the consequence of the detachment of these regions from the nuclear periphery (Zheng et al., 2018). The nuclear lamina being a nuclear compartment where several RIPPOs localise and where we could reconstitute H3K9me3 levels despite the absence of RIF1, suggests that this might be a domain characterised by high PP1 activity, crucial for maintaining and transmitting heterochromatin identity throughout the cell cycle (Holla et al., 2020): in our tethering experiment we have possibly channelled a normally RIF1-dependent region into a peripheral domain where another RIPPO can provide the PP1 activity required for H3K9me3 inheritance.

## Author contributions

N.C. conducted the majority of the experiments; R.R.N.P.K. helped with libraries preparation, ChIP and ELISA; B.P. wrote the pipeline and helped with the analysis for Fig. 5F and G; D.S. initiated the creation of the cell lines for targeting the TetO; S.B. performed the iPOND and prepare the proteins for the mass spectrometry experiments; S.W. performed the analysis of the ChIP-seq data; T.S. performed the replication timing experiments and analysis; SdS and M.V-A. have performed the histone mass spectrometry; S.C.B.B. conceived the project and wrote the manuscript with N.C; all the authors have provided feedback.

## Supporting information

all additional information

## Acknowledgements

We are grateful to the Bickmore lab (University of Edinburgh-UoE) for providing the plasmids containing the TetO array and the plasmid encoding for the mini-Lap2b cDNA. Shelag Boyd in the Bickmore lab and Rafael Czapiewski from the Schirmer Lab (UoE) helped with the 3D-FISH; Dirk Kleinjan, from the Rosser Lab (UoE) provided the plasmids for TetO array targeting and the Pollard lab (UoE) the plasmids to target the OsTir1 in the *Rosa26* locus. The Shimura lab is thanked for sharing the monoclonal antibodies against H3S10ph, CMC313 and CMC311; David Kelly from the COIL microscopy facility at the UoE is thanked for assistance. We also thank Martin Waterfall from the IIIR Flow Cytometry Core Facility at the UoE. We thank Caitlin Reid and Kumiko Samejima, from the Earnshaw group, for training and use of the elutriator and the cytospin.

