## Supplementary material for "RIF1 orchestrates a multi-step restoration of post-replicative H3K9me3": all additional information

#### Supplemental Data

**Supplemental Figure Legends**

**Supplemental Figures**

**Additional data figures**

**Materials and Methods**

**Table 1**

**Table 2**

#### Supplemental Figures

**Supplemental Figure 1. In Lamin B1-intermediate regions (cluster 2) peripheral localisation and B compartment association are RIF1-dependent. A.** Scatter plot of RIF1 enrichment (ChIP) versus Lamin B1 levels (DamID). Each dot represents a TAD. The regions analysed by 3D-FISH in D are shown. **B.** Replication timing (RT; E=early replicating; L=late replicating) over the genomic clusters defined in Fig.1A. Rif1-WT and Rif1-KO are shown. The values indicating early replication are shadowed in green, late in red. RT values were averaged across 5 kb windows. **C.** Genomic tracks of selected regions L9, O6 and L5. Lamin B1 DamID (Peric-Hupkes et al., 2010) and RIF1 ChIP (Foti et al., 2016) from wild type ESCs; replication timing (Foti et al., 2016) and chromatin compartments (eigenvector, (Gnan et al., 2021) from Rif1-WT and Rif1-KO cells are shown. A=A compartment, or positive eigenvector values; B= B compartment, or negative eigenvector values. The regions where the FISH probes hybridise are shadowed. TADs' boundaries are indicated by the lines on top. **D.** IF-3D-FISH analysis of additional cluster 2 regions**.** Lamina-association was evaluated as described in Fig.1C and Materials and Methods. Each point represents an independent Rif1-WT or Rif1-KO clone. The mean and standard deviation are shown in the plot. The statistical analysis was performed with Welch's *t*-test, $p<0.05$*. $p<0.01$**, $p<0.001$***.

**Supplemental Figure 2. The peripheral localisation and association with the B compartment of Lamin B1 intermediate genomic regions depends on the histone H3K9 methyltransferases SUV39H1/2. A.** Box plot of H3K9me3 ChIP log2(IP/input) across three RIF1-high clusters in Suv39h1/2 dn and control wild type cells. Data from (Bulut-Karslioglu et al., 2014). The H3K9me3 signal was averaged over TADs. The median and the interquartile range (± 1.5XIQR) are annotated. The statistical analysis was performed by Mann-Whitney U test**,** $p<0.05$*. $p<0.01$**, $p<0.001$***. **B.** A and B compartment analysis across the genomic clusters defined in Fig. 1A. in Suv39h1/2 dn and control wild type ESCs. The box plot summarises the distribution of the eigenvectors values. The A compartment is shadowed in green, the B in red. The statistical analysis was performed by Mann-Whitney U test, $p<0.05$*. $p<0.01$**, $p<0.001$***. Data from (Fukuda et al., 2021). **C.** The subnuclear localisation of the indicated genomic regions was analysed by IF-3D-FISH as in Fig.1C. The percentage of nuclei showing at least one lamina-associated signal was plotted. The mean and standard deviation of three independent experiments are shown. *p*-values were calculated by Welch's *t*-test, $p<0.05$*. $p<0.01$**, $p<0.001$***. **D.** RIF1 (Foti et al., 2016) and Lamin B1 (Peric-Hupkes et al., 2010) levels from wild type ESCs; H3K9me3 levels (ChIP-(Bulut-Karslioglu et al., 2014) and compartments (eigenvectors-(Fukuda et al., 2021) from Suv39h1/2 dn and control wild type ESCs are shown for the indicated regions, analysed by IF-3D-FISH in C. The regions recognised by the FISH probes are shadowed. A=A compartment, or positive eigenvector values; B= B compartment, or negative eigenvector values.

**Supplemental Figure 3: Loss of H3K9me3 at RIF1-bound loci**. **A.** Comparative analysis of H3K9me3 ChIP signals over the 3 RIF1-low clusters (clusters 4,5 and 6) between Rif1-WT and Rif1-KO. **B.** Genomic tracks of H3K9me3 from cluster 1 and 2 example regions in individual ESC clones. The L9 and O6 genomic regions shown are the same regions also targeted by the FISH probes used to analyse subnuclear positioning. The regions where the FISH probes hybridise are shadowed in grey; in yellow are highlighted some of the differential H3K9me3 peaks. Lamin B1 DamID and RIF1 ChIP are also displayed. **C**. ERVKs selected for H3K9me3 log2(H3K9me3_WT/Input) >0.1 are enriched in genomic regions falling within cluster 1. To assess the significance of ERVK co-localisation with these genomic domains, permutated sequences matching the size of ERVKs were analysed. The *p*-values were calculated by permutation test. **D.** Analysis of RIF1 ChIP-seq (Foti et al., 2016) reveals that RIF1 peaks are enriched in all ERV categories, as defined in A. ERVKs show the strongest enrichment. The *p*-values were calculated by permutation test. **E.** RIF1 is significantly enriched over the ERVs that display reduced H3K9me3 in Rif1-KO cells, as shown in Fig. 2B. The *p*-values were calculated by permutation test. **F.** The reduction of H3K9me3 in Rif1-KO cells significantly coincides with RIF1 binding sites across the different clusters. The *p*-values were calculated by permutation test.

**Supplemental Figure 4. Tethering to the nuclear periphery and restoring H3K9me3 levels is not sufficient to reinstate late replication timing of a cluster 2 genomic region. A.** Illustration of the tethering platform. *TetO* arrays were integrated in the O2 region (cluster 2, see Material and Methods). The anchor is a fusion protein FRB-mini-Lap2$\beta$, located to the nuclear periphery; the tether is TetR-FKBP. The tethering of O6-TetO to the nuclear periphery is achieved by Rapamycin-induced dimerization of FRB and

FKBP. **B.** Dual colour FISH with a probe against the *TetO* array (green) and a second probe against the O6 genomic region (red), showing that the *TetO* array is integrated into one of the two O6 alleles (see Additional Information Fig.4). Scale bars=3μm. **C.** Schematic of the construct driving the expression of the anchor and the tether (top). **D.** Top: Immunostaining with anti-Myc antibody demonstrates that the Myc-FRB-miniLap2β fusion protein is associated with the nuclear periphery (green), as shown by the overlap with Lamin B1 (red). Middle: Immunostaining with anti-HA (green) detects HA-TetR-FKBP throughout the nucleoplasm in cells treated with DMSO. The signal becomes concentrated at the lamina (Lamina B1, red) upon treatment with Rapamycin, indicating that the dimerisation between FRB and FKPB brings HA-TetR-FKBP to the lamina. Bottom: in DMSO-treated cells pre-extracted with Triton X-100 buffer prior to fixation, the anti-HA signal is removed everywhere but in a focus, presumably corresponding to the *TetO*, where the HA-TetR-FKBP is retained as it is tightly bound to DNA. After treatment with Rapamycin, the signal is concentrated at the periphery. Scale bars=3μm. **E.** Experimental design to tether O6 back to the nuclear periphery in the absence of RIF1: over a 4 days OHT treatment course to delete *Rif1* (see Materials and Methods), Rapamycin was added during the last 2 days. **F.** The efficiency of Rapamycin-induced tethering of the TetO-containing O6 region was evaluated by Lamin B1 IF-3D-FISH. The mean and standard deviation of three independent experiments is shown in the plot. The treatment with Rapamycin increases the association of O6 with the nuclear lamina also in the control cells, pushing it towards the higher frequencies that are typical of cluster 1 regions. The statistical analysis was performed by Welch's *t*-test, $p<0.05$*. $p<0.01$**, $p<0.001$***. **G.** ChIP-qPCR analysis of H3K9me3 levels at the tethered O6-TetO region, for the conditions shown in E. Two different primer pairs were used, one specific for the inserted *TetO* array (Attb-TetO), the other recognising both alleles of O6. The levels of H3K9me3 at the indicated regions were normalised against the internal control β-actin. **H.** Replication timing profiles of cells in the presence or absence of RIF1 (WT and KO respectively) with DMSO or Rapamycin treatment in three independent experiments. The O6 region targeted by the FISH probe is shadowed, and the *TetO* integration site is highlighted by the dashed line.

**Supplemental Figure 5. RIF1 associates with chromatin during mitotic exit. Acute depletion of RIF1 via degron. A.** A cell line where a mCherry tag was knocked-in at the

N-terminus of *Rif1* was fixed and imaged. H3S10ph was used as a marker of mitotic progression. The white arrows mark telophase cells where H3S10ph signal is normally strongly reduced or completely removed. Scale bar=3μm. **B.** Schematic of the N-terminal knock-in of an auxin-inducible degron (mini-AID-mCherry) tag in the *Rif1* locus. **C.** Treatment of $^{\text{mini-AID-mCherrry/mini-AID-}\textit{mCherrry}}$*Rif1* cells with auxin-analogue 5-ph-IAA for 2 hrs induces RIF1 degradation (red). Arrows highlight telophase cells where H3S10ph (green) should not be detectable anymore (see A): after inducing RIF1 degradation, H3S10ph dephosphorylation is delayed and the signal is still detectable is some telophase cells. **D.** Western blot probed with anti-mouse RIF1 antibody 1240 and anti-SMC1 as loading control. The indicated different concentration of 5-ph-IAA were used for the indicated different times to induce acute degradation of RIF1.

**Supplemental Figure 6. RIF1 depletion specifically affects the levels of H3K9me3. A.** Flow cytometry shows a comparable efficiency of release from nocodazole arrest between DMSO and 5-ph-IAA-treated cells. One representative experiment is shown. G1 gate: cells that were pulsed with EdU during Mid-Late S phase (as shown in Fig.3A) enter the next G1 phase after being released from Nocodazole for 2 hrs. G2 gate: EdU pulsed cells that remain in G2/M after nocodazole release. **B.** Flow cytometry analysis on PFA fixed and DAPI stained cells shows the reduction of mCherry signal (RIF1) in 5-ph-IAA-treated cells released from nocodazole. One representative experiment is shown. **C.** The efficiency of 5-ph-IAA-induced RIF1 degradation evaluated by western blot in two independent experiments. RIF1 was visualised using a Rabbit polyclonal in-house anti-mouse antibody (1240); anti-SMC1 antibody was used for the loading control. **D.** Barplot summarising the proportion of incorporation of old (light) and newly synthesised (heavy) H3 and H4 histones versus the total histones in DMSO and 5-ph-IAA treated cells. Error bars indicate standard deviations. DMSO (n=2), IAA (n=3). **E.** Barplot summarising the relative abundance of H3K9me3_K14 and H3K9me2_K14. DMSO (n=2), IAA (n=3). The mean and standard deviation are shown. *p*-values were calculated using a Welch's *t*-test. **F.** Heatmap of the average percentage (DMSO n=2; IAA n=3) of indicated modifications (H3K9me0/1/2/3/ac_K14ac) over the total histone H3 peptide aminoacids (aa) 9-17. Both parental (light) or newly incorporated (heavy) histone H3 extracted from G1 cells were analysed. **G.** Barplot shows the percentage of H3K9me2/3_K14ac relative to the total histone H3 aa 9-17. DMSO (n=2), IAA (n=3). The mean and standard deviation are shown. Statistical analysis was performed by a Welch's *t*-test. **H.** Barplot shows the ratio of the relative abundance of H3K9me3_K14ac and H3K9me2_K14ac. DMSO (n=2), IAA (n=3). The mean and standard deviation are shown. Statistical analysis was performed by a Welch's *t*-test. **I.** Barplot shows the percentage of H3.1K27me2/3 (top) and H3.1K36me2/3 (bottom) relative to the total histone H3.1 aa 27-40. DMSO (n=2), IAA (n=3). The mean and standard deviation are shown. Statistical analysis was performed by a Welch's *t*-test. **J.** Heatmap of the relative abundance of identified peptides on old (light) or new (heavy) histone H3.1 peptide aa 27-40 in G1 cells, indicated as the average of percentage of enrichment (DMSO n=2; IAA n=3).

**Supplemental Figure 7. RIF1 depletion does not affect the total levels of proteins involved in H3K9me3 restoration. A.** Western blots quantifying the total level of several proteins involved in H3K9me3 restoration, some of which are found specifically reduced on mid-late replicating chromatin isolated in the following G1 in Fig. 3E. Results from two representative experiments are shown. **B.** PCA analysis summarising the results from the 4 independent experiments that were analysed by iPOND-MS.

**Supplemental Figure 8. RIF1-PP1 is important for the maintenance of H3K9me3 levels. A.** Heatmap of H3K9me3 Z-scores assessed across ERVs pre-selected for a minimum normalised read count of 10 and with a log2(WT/Input) >0.1. The analysis includes results from three Rif1-WT, four Rif1-KO clones, three Rif1-HEMI and three Rif1-ΔPP1. **B.** Comparative analysis of H3K9me3 ChIP signals over the 3 RIF1-low clusters (clusters 4,5 and 6) between Rif1-HEMI and Rif1-ΔPP1. The signal was averaged over 5 kb windows. **C.** Genomic tracks of H3K9me3 in individual clones from regions L9 and O6. In grey are shadowed regions where the FISH probes hybridise; in yellow are highlighted some of the differential H3K9me3 peaks. Lamin B1 DamID and RIF1 ChIP are also displayed. **D.** IF-3D-FISH analysis of O6, L5 and L9 regions in Rif1-WT, Rif1-KO, Rif1-HEMI and Rif1-ΔPP1 cells. The mean and standard deviation of three independent experiments is shown in the plot. The statistical analysis was performed with Welch's *t*-test, p<0.05*. p<0.01**, p<0.001***. **E.** Eigenvector of genomic regions in three RIF1-high clusters (1, 2 and 3) fromRif1-WT, Rif1-KO, Rif1-HEMI and Rif1-ΔPP1 cells (Foti et al., 2016).

**Supplemental Figure 9. H3S10 is phosphorylated prematurely, before replication, in the absence of functional RIF1. A.** The levels of H3S10ph were quantified by flow cytometry in cycling $^{\text{mCherrry/mini-AID-}\textit{mCherrry}}$*Rif1* cells after 4 days of 5-ph-IAA treatment. DNA was quantified by DAPI staining. The DAPI signal of all conditions was normalised to the respective G1 cells. G1/early S cells were defined by a normalised DAPI signal < 1.3; early/mid S phase cells were defined by normalised DAPI signal between 1.3-1.5. Hesperadin treatments for 2 hrs prior to collecting cells reverts the gain of H3S10ph in RIF1-depleted cells. Top FACS profile with gates; bottom, quantification of the H3S10ph signal in the indicated DAPI intervals. **B.** Analysis of Aurora B levels in elutriated G1 cells, across Rif1-WT, Rif1-KO, Rif1-ΔPP1 and Rif1-HEMI treated with OHT for four days. Top: FACS profile of elutriated cells stained by DAPI. Bottom: western blot. The total levels of Aurora B in G1 and in cycling cells (Pre) appear comparable across the different cell lines. Histone H3 was used as a loading control. **C**. $^{\text{mCherrry/mini-AID-}\textit{mCherrry}}$*Rif1* cells were synchronised and released, in presence (DMSO) or absence (5-ph-IAA) of RIF1 during mitosis, as described in Fig.3A, in normal media. Cells released from nocodazole for 2 hrs were collected and fixed with PFA and then stained with Propidium iodide (PI) for 2 hrs prior to sorting. Aurora B levels were analysed by western blot. Left: exemplary FACS profile of PI stained, sorted cells. Right: western blot probed with anti-Aurora B antibody and anti-histone H3 as a loading control. **D.** Immunostaining for H3S10ph (Millipore antibody 06-570) and MCM3 in cycling Rif1-WT, Rif1-KO and Rif1-ΔPP1 cells. EdU staining was used as a reference for the position of the replication forks and the S-phase substage. MCM3 binds to chromocenters yet to be replicated (EdU negative) or replicating (EdU positive). In wild type cells, MCM3 and H3S10ph do not co-localise, as H3S10ph only binds chromocenters after replication (see insets). On the contrary, MCM3 and H3S10ph overlap on both EdU negative and EdU positive chromocenters in Rif1-KO and Rif1-ΔPP1 cells. **E**. Same as in D, with a different anti-H3S10ph antibody, the monoclonal CMA313 (Hayashi-Takanaka et al., 2009).

Supp. Fig.1

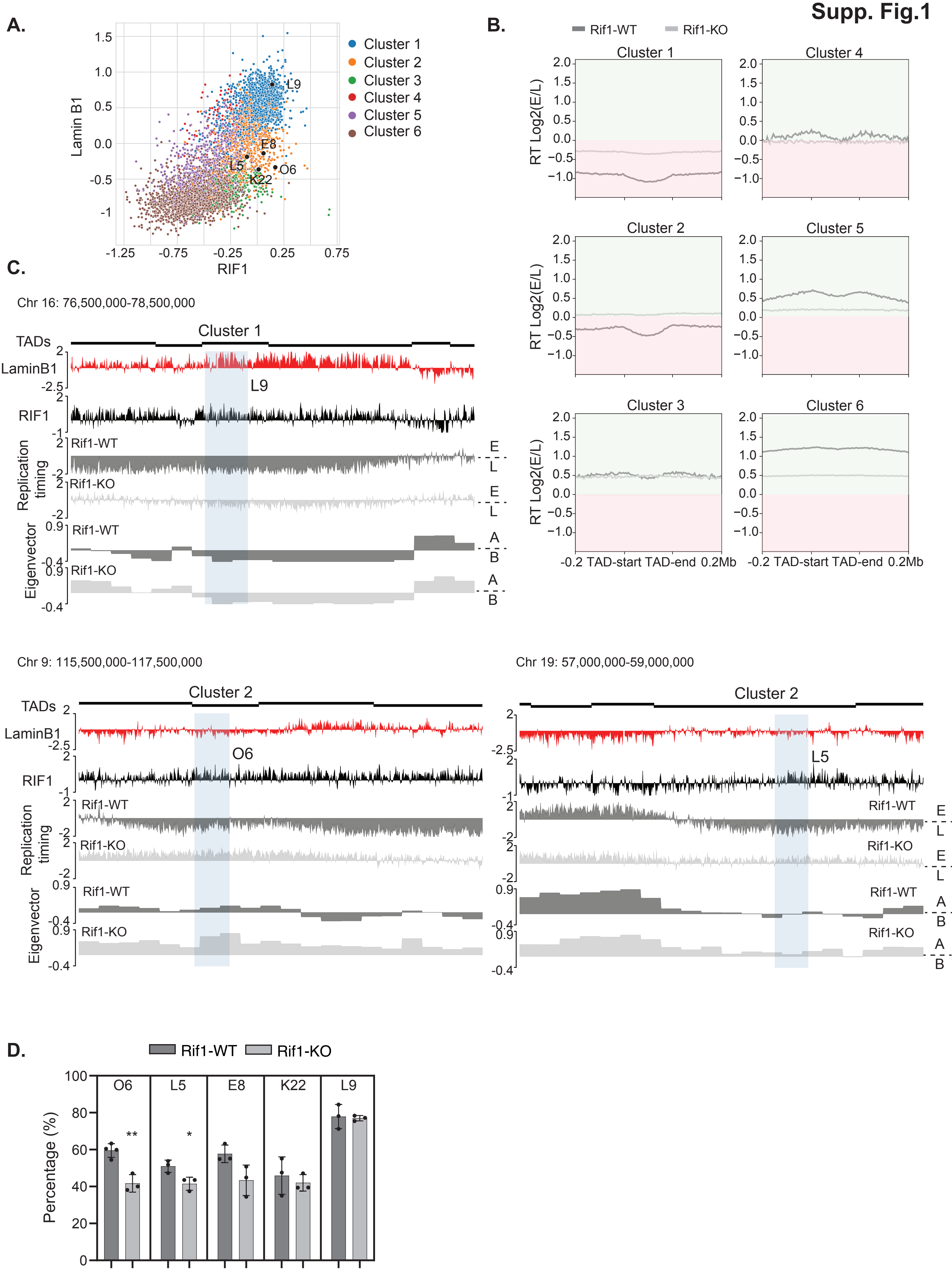
A.
Lamin B1
RIF1
Cluster 1
Cluster 2
Cluster 3
Cluster 4
Cluster 5
Cluster 6
L9
E8
L5
O6
K22
B.
Rif1-WT
Rif1-KO
Cluster 1
Cluster 2
Cluster 3
Cluster 4
Cluster 5
Cluster 6
RT Log2(E/L)
-0.2 TAD-start TAD-end 0.2Mb
C.
Chr 16: 76,500,000-78,500,000
Cluster 1
TADs
LaminB1
RIF1
Replication timing
Eigenvector
Rif1-WT
Rif1-KO
E
L
A
B
L9
Chr 9: 115,500,000-117,500,000
Cluster 2
O6
Chr 19: 57,000,000-59,000,000
Cluster 2
L5
D.
Rif1-WT
Rif1-KO
Percentage (%)
O6
L5
E8
K22
L9
**
*

#### Supp. Fig.2

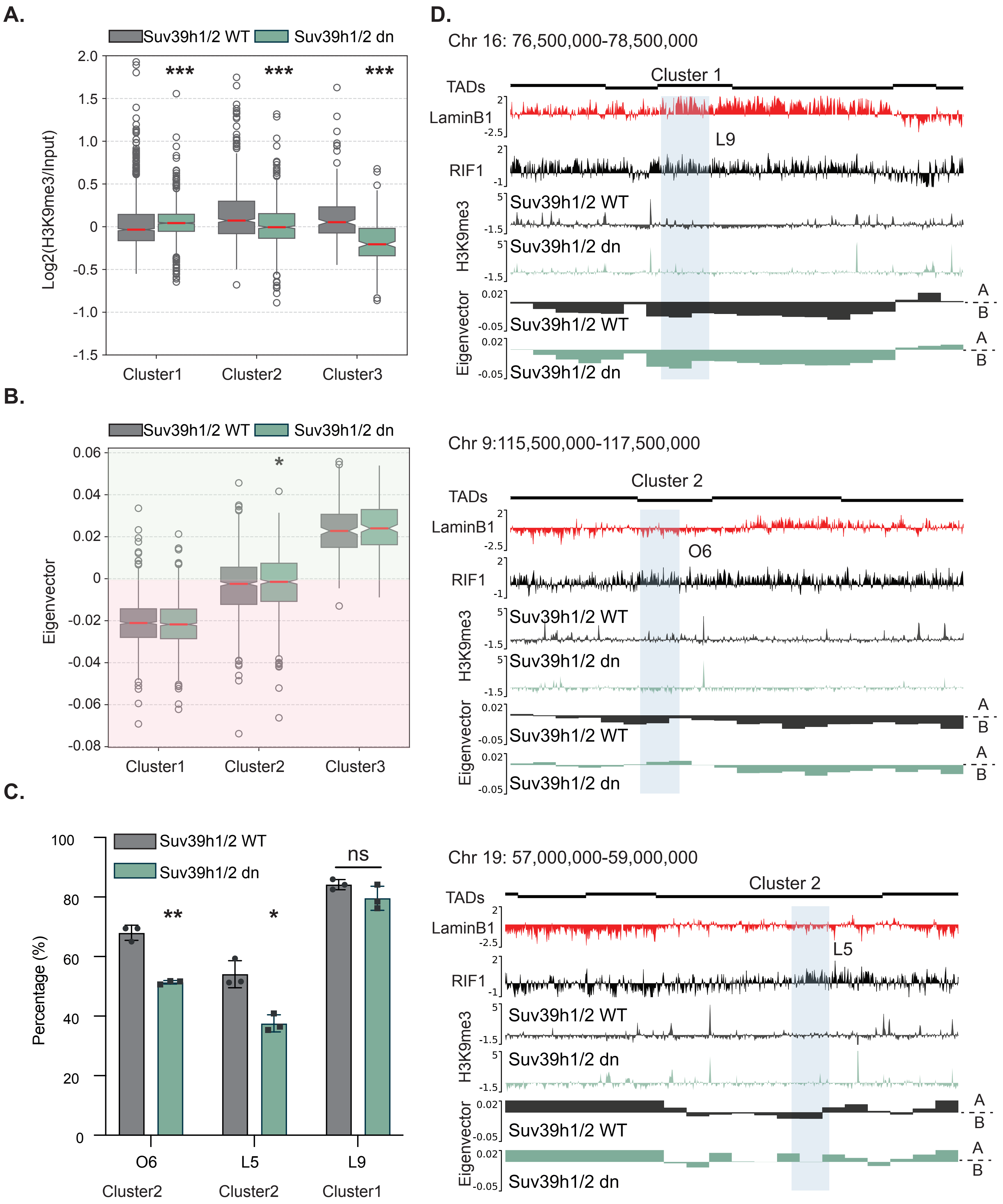

A.
Suv39h1/2 WT
Suv39h1/2 dn
Log2(H3K9me3/Input)
***
Cluster1
Cluster2
Cluster3
B.
Eigenvector
*
C.
Percentage (%)
ns
**
*
O6
L5
L9
D.
Chr 16: 76,500,000-78,500,000
Cluster 1
TADs
LaminB1
RIF1
H3K9me3
Eigenvector
A
B
Chr 9:115,500,000-117,500,000
Cluster 2
Chr 19: 57,000,000-59,000,000

**Supp. Fig.3**

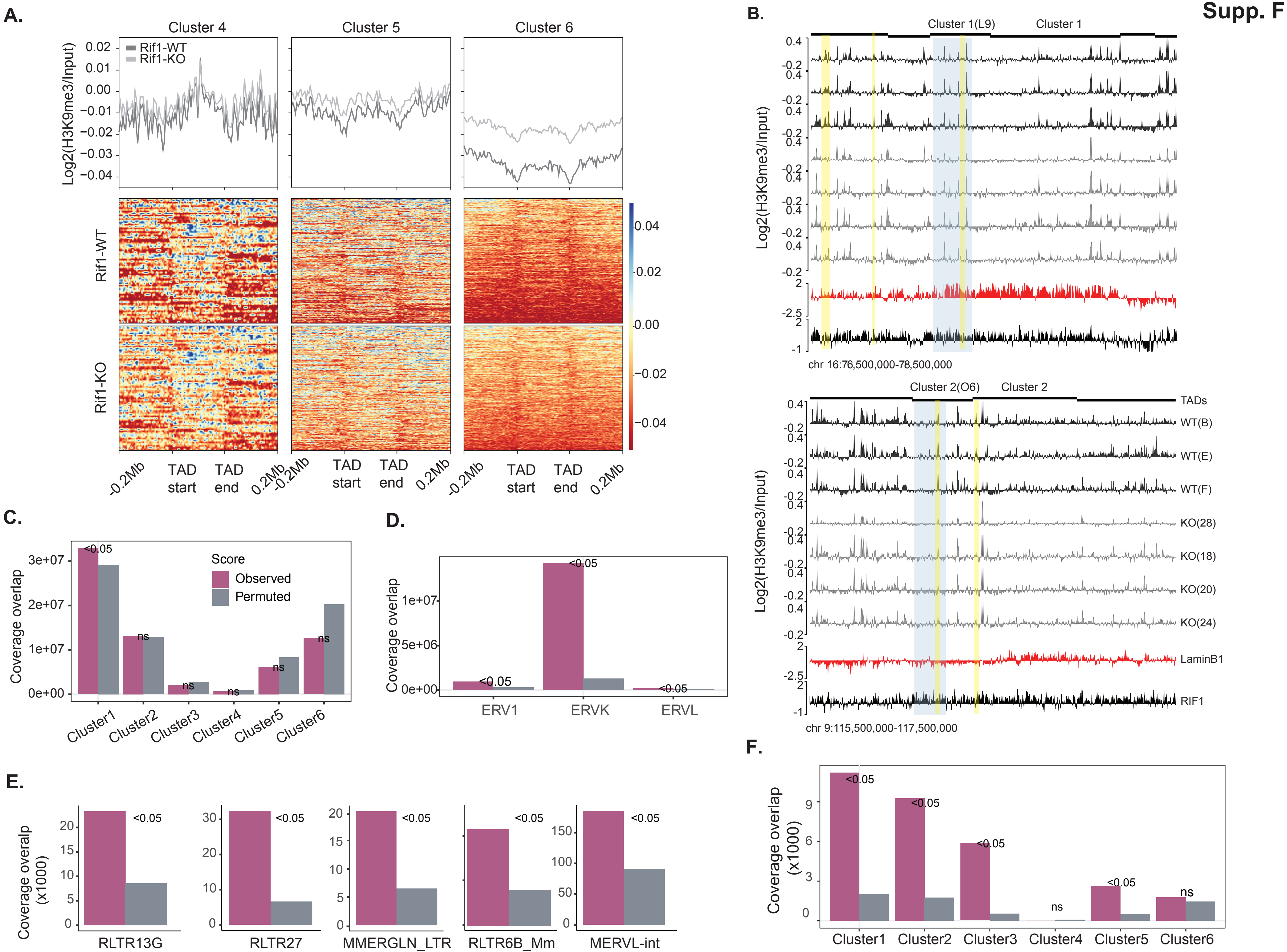
A.
Cluster 4
Cluster 5
Cluster 6
Rif1-WT
Rif1-KO
Log2(H3K9me3/Input)
Rif1-WT
Rif1-KO
-0.2Mb
TAD start
TAD end
0.2Mb
B.
Cluster 1(L9)
Cluster 1
chr 16:76,500,000-78,500,000
Cluster 2(O6)
Cluster 2
TADs
WT(B)
WT(E)
WT(F)
KO(28)
KO(18)
KO(20)
KO(24)
LaminB1
RIF1
chr 9:115,500,000-117,500,000
C.
Coverage overlap
Score
Observed
Permuted
Cluster1
Cluster2
Cluster3
Cluster4
Cluster5
Cluster6
D.
ERV1
ERVK
ERVL
E.
Coverage overalp (x1000)
RLTR13G
RLTR27
MMERGLN_LTR
RLTR6B_Mm
MERVL-int
F.
Coverage overlap (x1000)

#### Supp. Fig.4

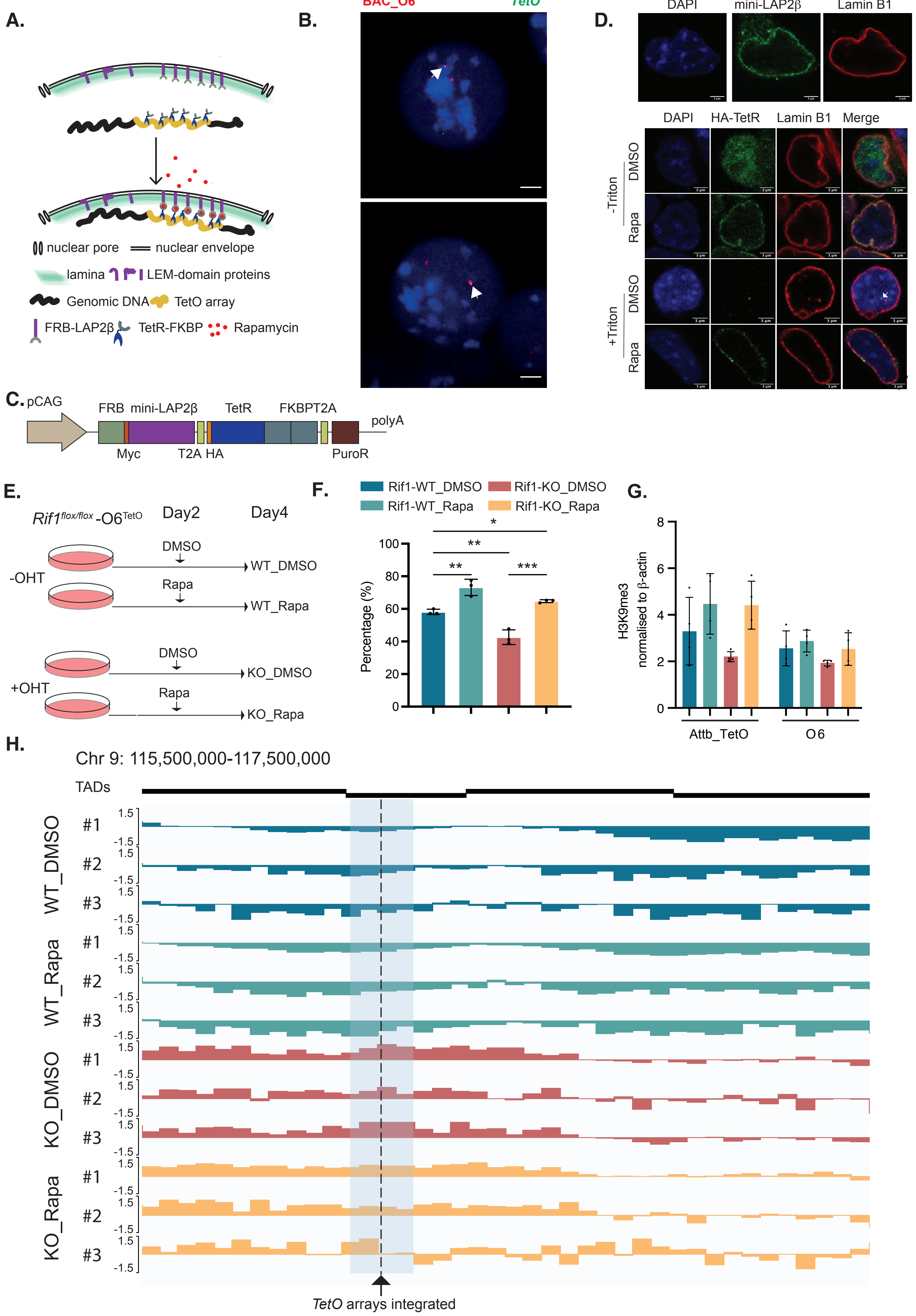

A.
nuclear pore
nuclear envelope
lamina
LEM-domain proteins
Genomic DNA
TetO array
FRB-LAP2β
TetR-FKBP
Rapamycin
B.
BAC_O6
TetO
C. pCAG
FRB mini-LAP2β
TetR
FKBPT2A
polyA
Myc
T2A HA
PuroR
D.
DAPI
mini-LAP2β
Lamin B1
DAPI
HA-TetR
Lamin B1
Merge
-Triton
+Triton
DMSO
Rapa
E.
Rif1 flox/flox -O6 TetO
Day2
Day4
-OHT
+OHT
DMSO
Rapa
WT_DMSO
WT_Rapa
KO_DMSO
KO_Rapa
F.
Rif1-WT_DMSO
Rif1-KO_DMSO
Rif1-WT_Rapa
Rif1-KO_Rapa
Percentage (%)
G.
H3K9me3 normalised to β-actin
Attb_TetO
O6
H.
Chr 9: 115,500,000-117,500,000
TADs
WT_DMSO
WT_Rapa
KO_DMSO
KO_Rapa
#1
#2
#3
TetO arrays integrated

Supp. Fig. 5

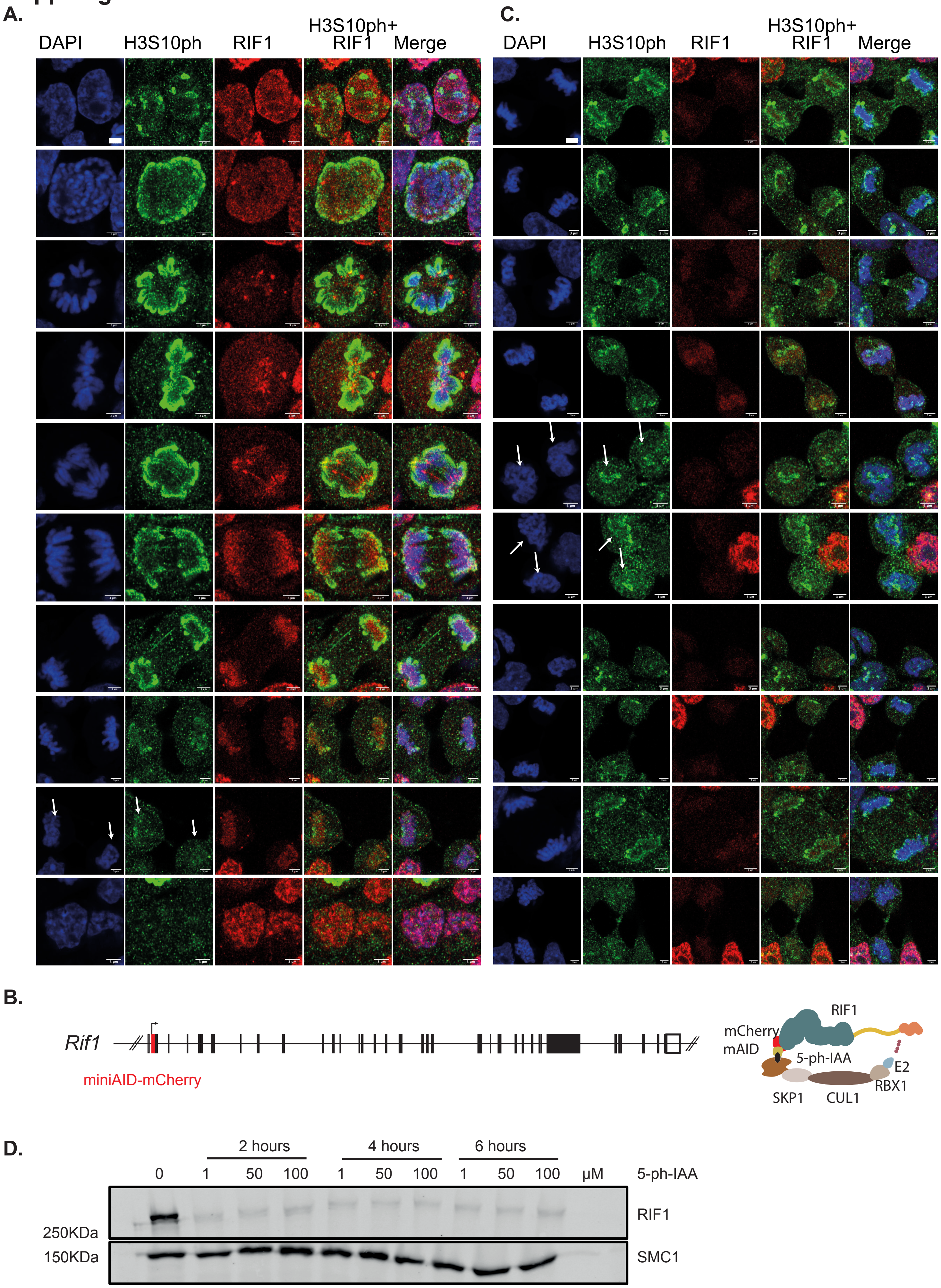

A.
DAPI
H3S10ph
RIF1
H3S10ph+ RIF1
Merge
C.
DAPI
H3S10ph
RIF1
H3S10ph+ RIF1
Merge
B.
Rif1
miniAID-mCherry
RIF1
mCherry
mAID
5-ph-IAA
E2
RBX1
SKP1
CUL1
D.
2 hours
4 hours
6 hours
0
1
50
100
1
50
100
1
50
100
μM
5-ph-IAA
RIF1
SMC1
250KDa
150KDa

#### Supp. Fig.6

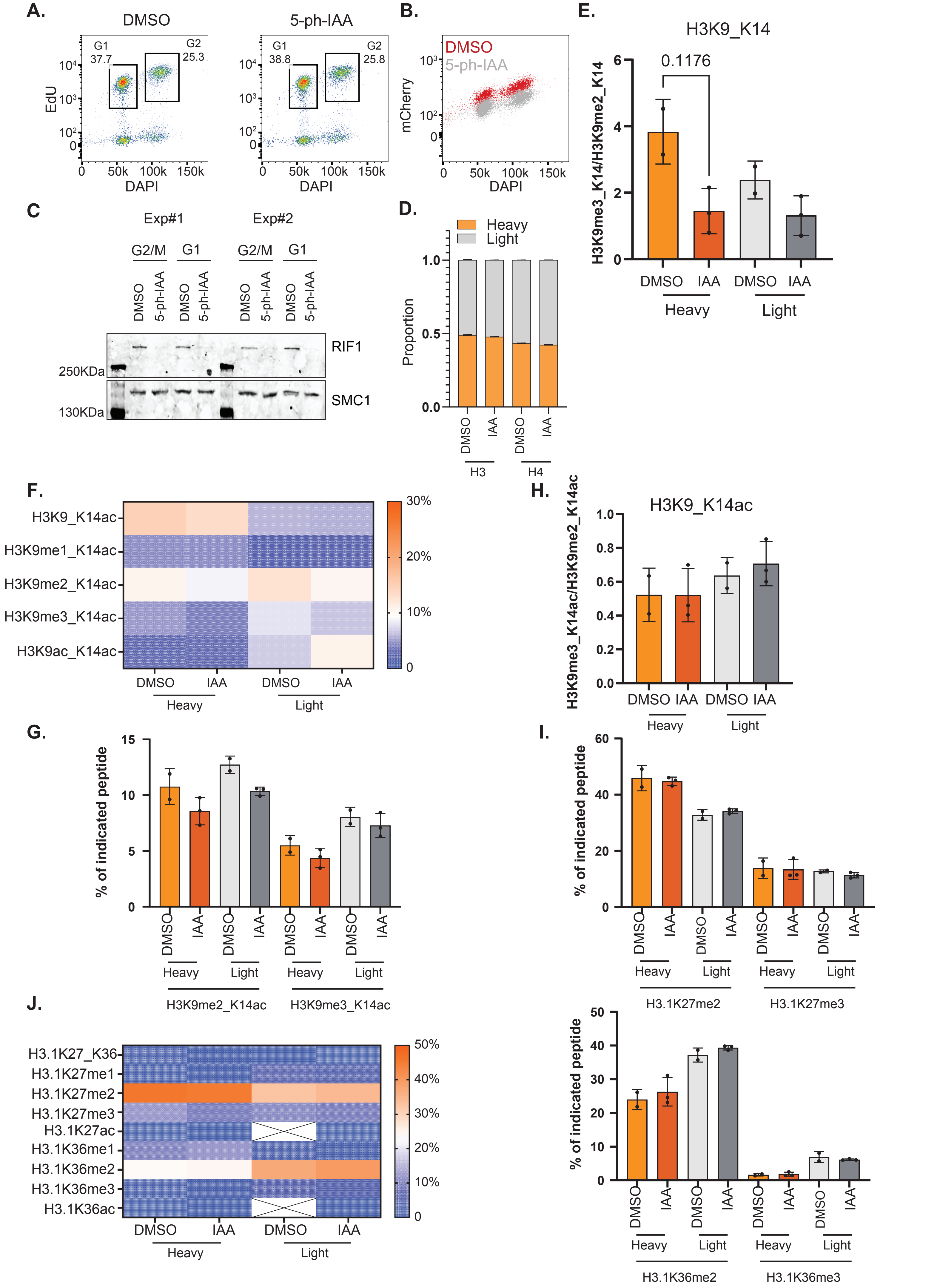

A.
DMSO
5-ph-IAA
G1 37.7
G2 25.3
G1 38.8
G2 25.8
EdU
DAPI
B.
DMSO
5-ph-IAA
mCherry
DAPI
C
Exp#1
Exp#2
G2/M
G1
DMSO
5-ph-IAA
RIF1
SMC1
250KDa
130KDa
D.
Heavy
Light
Proportion
H3
H4
E.
H3K9_K14
0.1176
H3K9me3_K14/H3K9me2_K14
DMSO
IAA
Heavy
Light
F.
H3K9_K14ac
H3K9me1_K14ac
H3K9me2_K14ac
H3K9me3_K14ac
H3K9ac_K14ac
30%
20%
10%
H.
H3K9_K14ac
H3K9me3_K14ac/H3K9me2_K14ac
G.
% of indicated peptide
H3K9me2_K14ac
H3K9me3_K14ac
I.
H3.1K27me2
H3.1K27me3
J.
H3.1K27_K36
H3.1K27me1
H3.1K27me2
H3.1K27me3
H3.1K27ac
H3.1K36me1
H3.1K36me2
H3.1K36me3
H3.1K36ac
50%
40%
30%
20%
10%
H3.1K36me2
H3.1K36me3

### Supp. Fig.7

A.

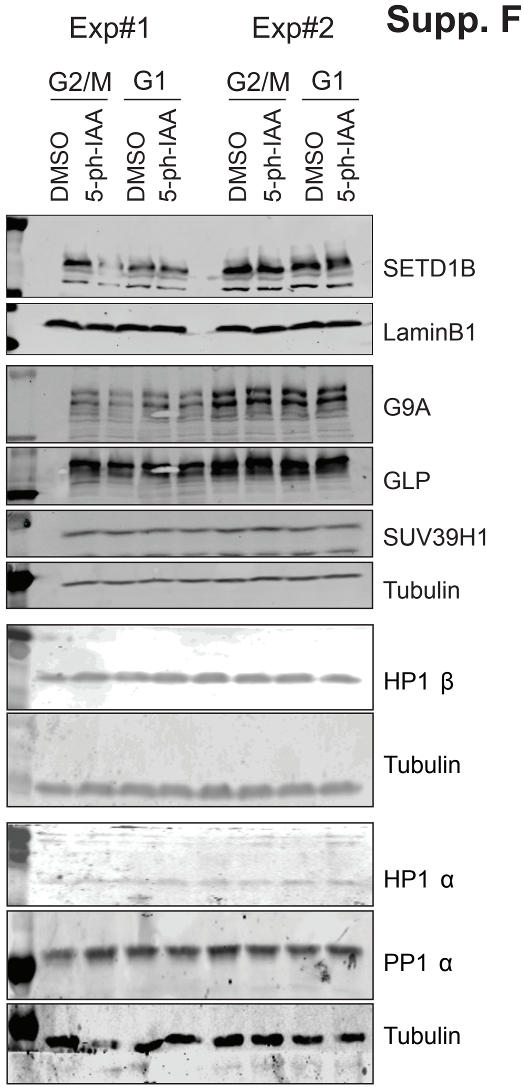

Exp#1
Exp#2
G2/M
G1
G2/M
G1
DMSO
5-ph-IAA
DMSO
5-ph-IAA
DMSO
5-ph-IAA
DMSO
5-ph-IAA
SETD1B
LaminB1
G9A
GLP
SUV39H1
Tubulin
HP1 β
Tubulin
HP1 α
PP1 α
Tubulin

B.

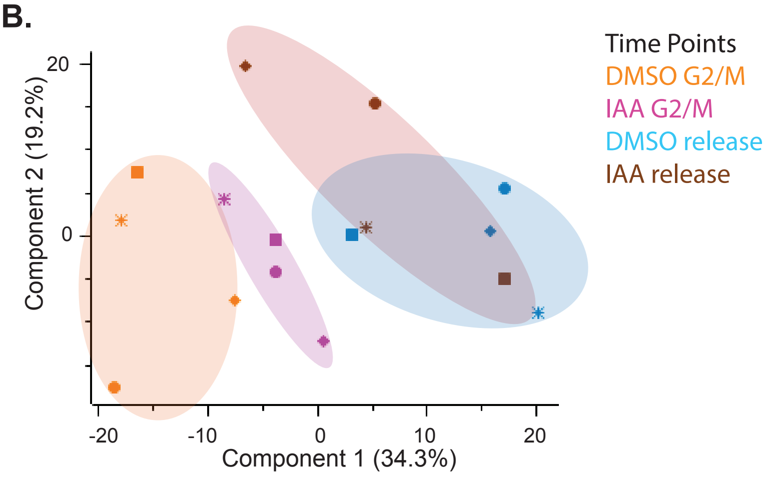

Component 2 (19.2%)
Component 1 (34.3%)
20
0
-20
-10
0
10
20
Time Points
DMSO G2/M
IAA G2/M
DMSO release
IAA release

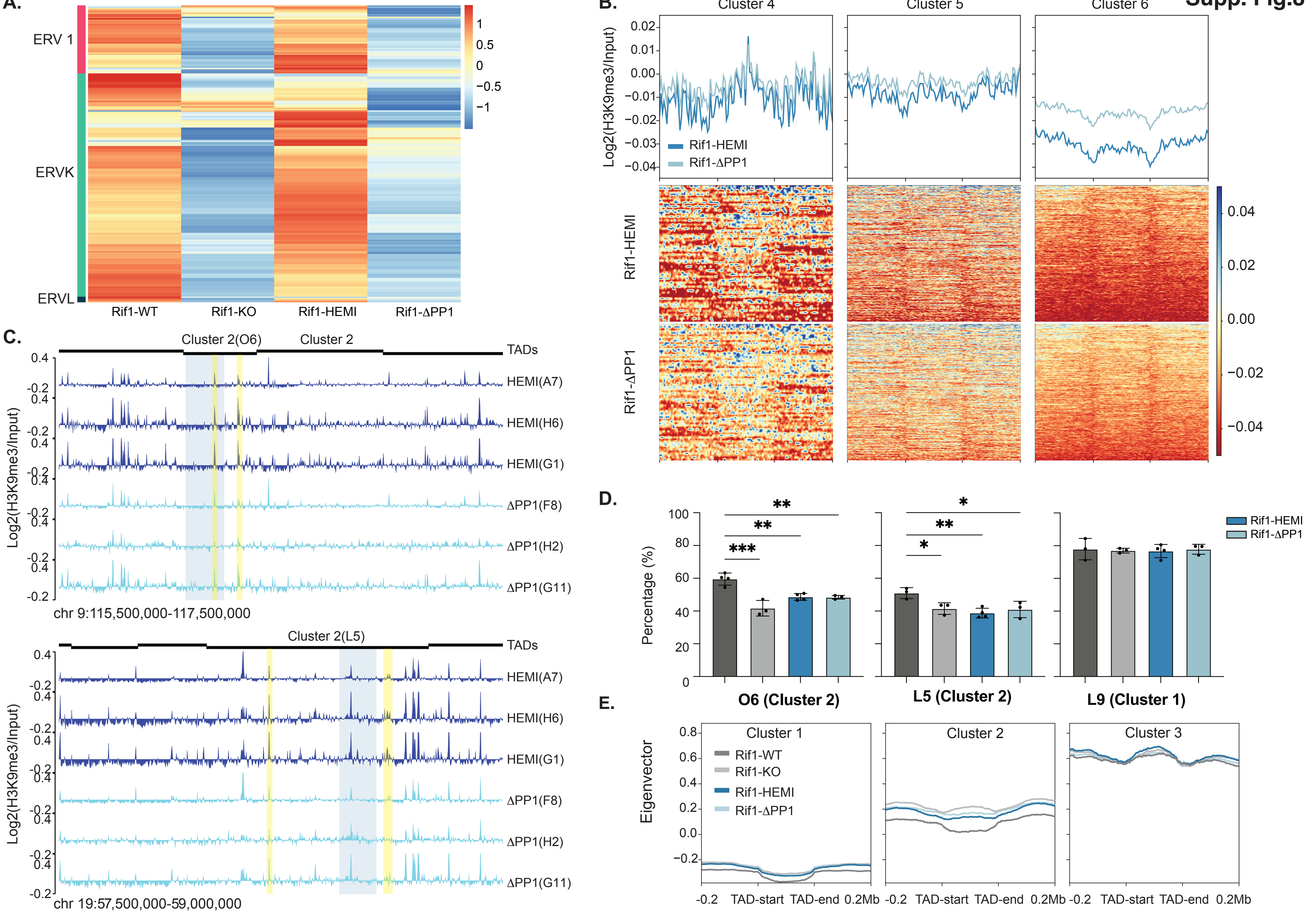
A.
ERV 1
ERVK
ERVL
Rif1-WT
Rif1-KO
Rif1-HEMI
Rif1-ΔPP1
1
0.5
0
−0.5
−1
B.
Cluster 4
Cluster 5
Cluster 6
Log2(H3K9me3/Input)
Rif1-HEMI
Rif1-ΔPP1
0.02
0.01
0.00
−0.01
−0.02
−0.03
−0.04
0.04
0.02
0.00
−0.02
−0.04
C.
Cluster 2(O6)
Cluster 2
TADs
HEMI(A7)
HEMI(H6)
HEMI(G1)
ΔPP1(F8)
ΔPP1(H2)
ΔPP1(G11)
Log2(H3K9me3/Input)
chr 9:115,500,000-117,500,000
Cluster 2(L5)
chr 19:57,500,000-59,000,000
D.
Percentage (%)
O6 (Cluster 2)
L5 (Cluster 2)
L9 (Cluster 1)
E.
Eigenvector
Cluster 1
Cluster 2
Cluster 3
Rif1-WT
Rif1-KO
Rif1-HEMI
Rif1-ΔPP1
-0.2
TAD-start
TAD-end
0.2Mb

Supp. Fig.9

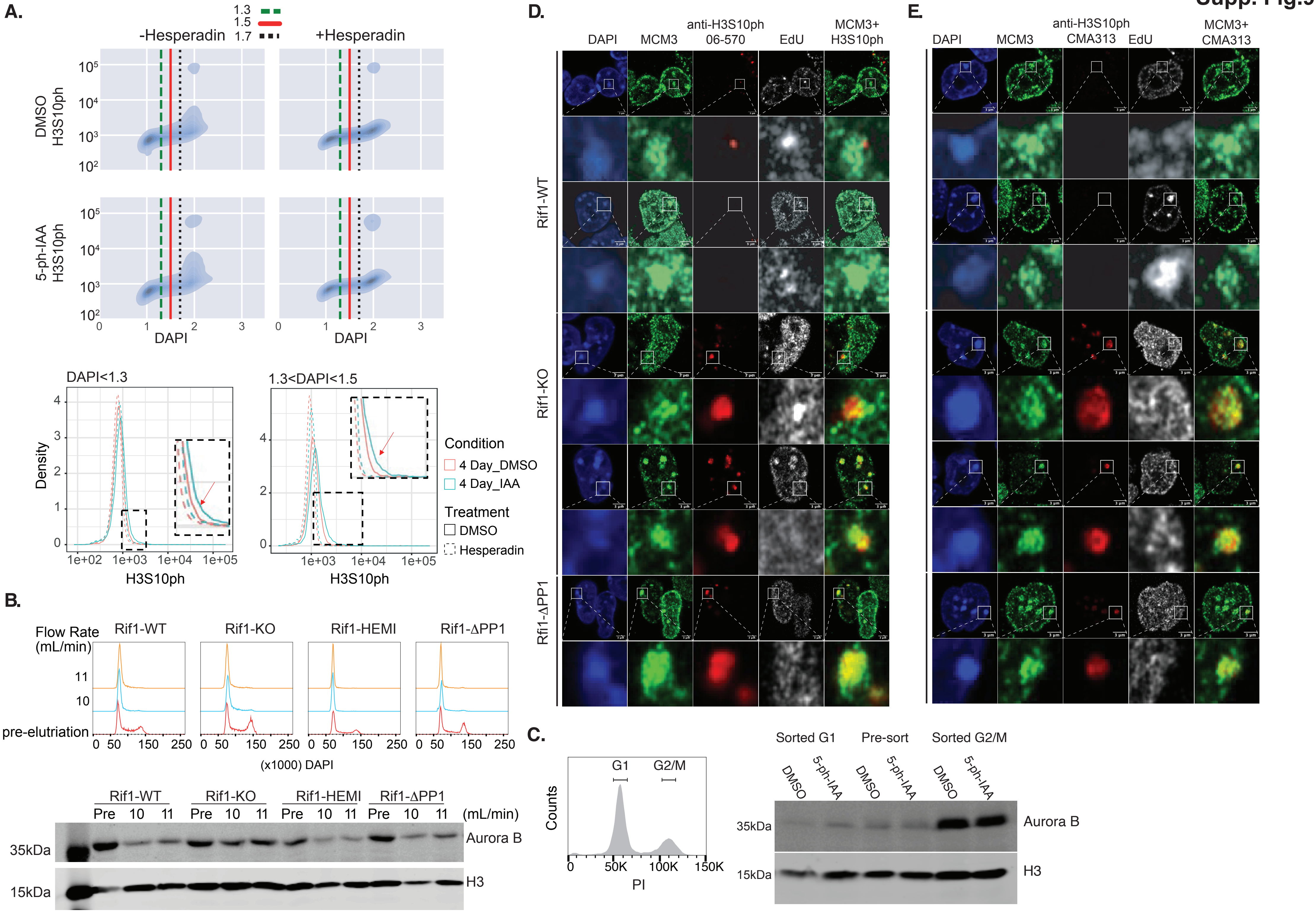
A.
1.3
1.5
1.7
-Hesperadin
+Hesperadin
DMSO
H3S10ph
5-ph-IAA
H3S10ph
$10^5$
$10^4$
$10^3$
$10^2$
0 1 2 3
DAPI
DAPI<1.3
1.3<DAPI<1.5
Density
H3S10ph
1e+02 1e+03 1e+04 1e+05
Condition
4 Day_DMSO
4 Day_IAA
Treatment
DMSO
Hesperadin
B.
Flow Rate
(mL/min)
Rif1-WT
Rif1-KO
Rif1-HEMI
Rif1-ΔPP1
11
10
pre-elutriation
0 50 150 250
(x1000) DAPI
Pre 10 11
(mL/min)
35kDa
15kDa
Aurora B
H3
C.
Counts
G1
G2/M
0 50K 100K 150K
PI
Sorted G1
Pre-sort
Sorted G2/M
DMSO
5-ph-IAA
35kDa
15kDa
Aurora B
H3
D.
DAPI
MCM3
anti-H3S10ph
06-570
EdU
MCM3+
H3S10ph
Rif1-WT
Rif1-KO
Rfi1-ΔPP1
E.
DAPI
MCM3
anti-H3S10ph
CMA313
EdU
MCM3+
CMA313

### Additional information Figure 1

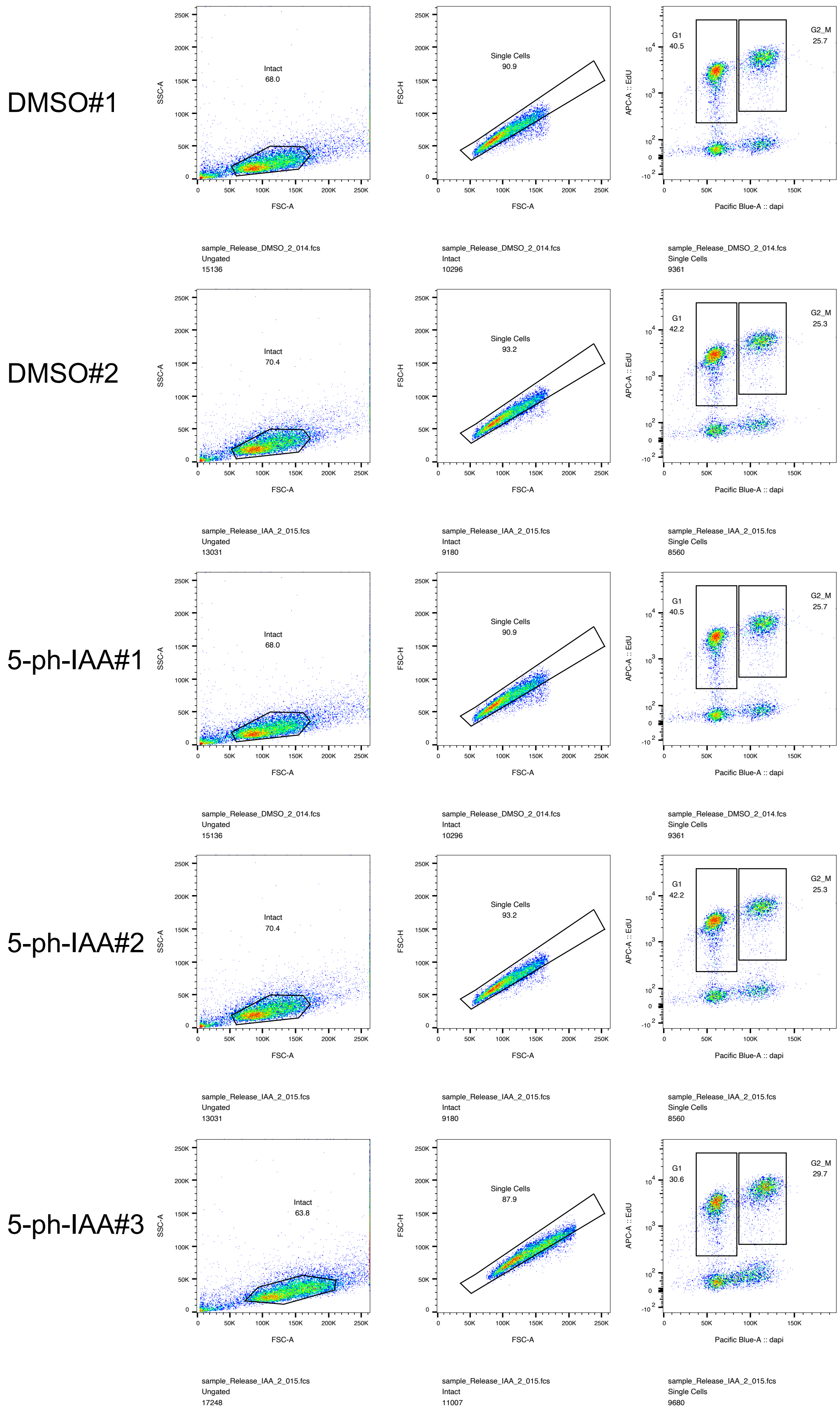

**Additional information Figure 1.** Flow cytometry analysis of samples used for SILAC-iPOND-MS in Figure 3. Cells synchronised and EdU labelled as in Figure 3A were collected after release from nocodazole. A fraction of cells was used to evaluate the release by flow cytometry. DMSO, n=2; 5-ph-IAA, n=3. The gating strategy is shown. EdU positive G1 (released from nocodazole) and G2_M are gated.

#### Additional information Figure 2

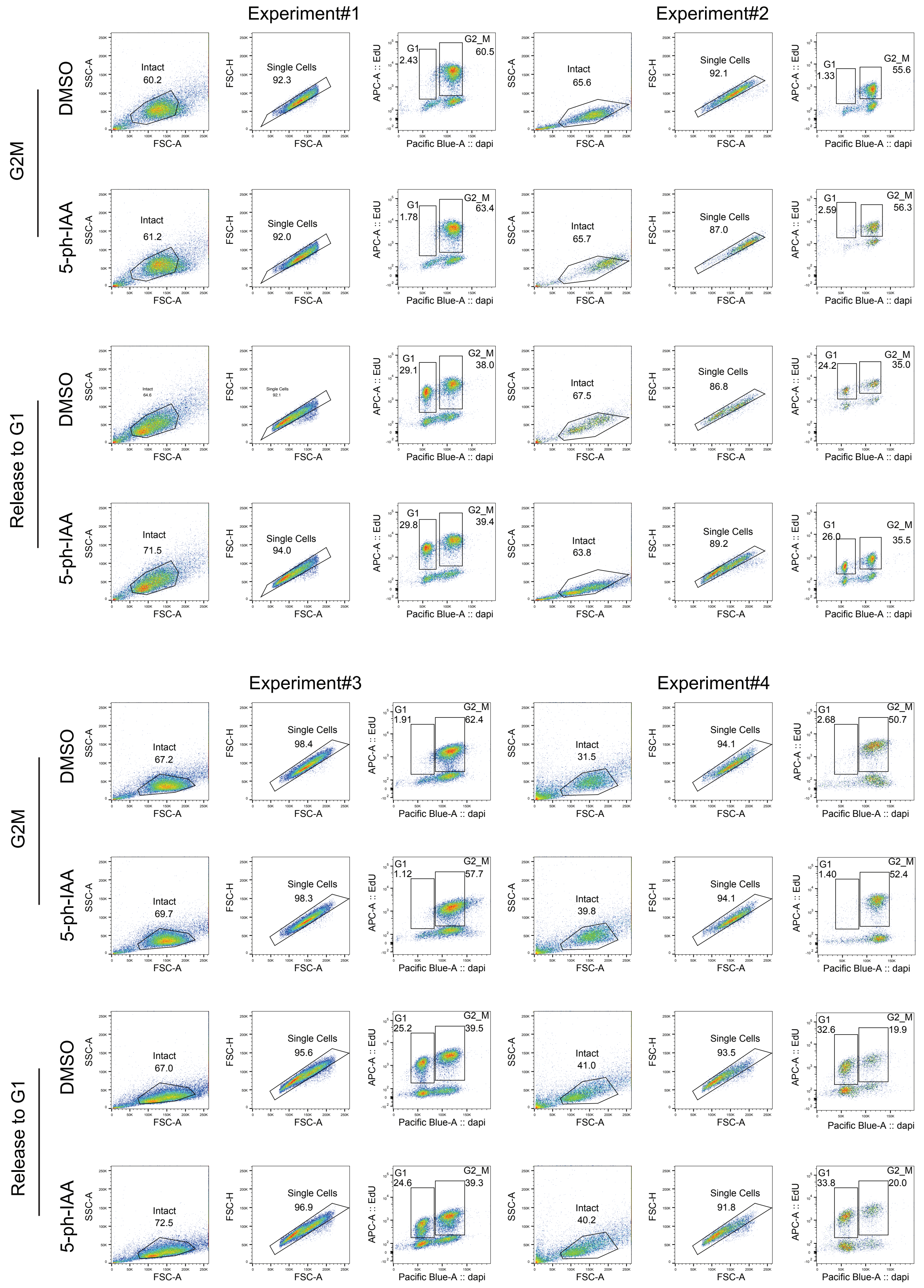

**Additional information Figure 2.**Flow cytometry analysis of samples used for iPOND-MS in Figure 3. The samples were prepared as described in the legend of Figure 3E. A fraction of the cells was callected to validate the release by flow cytometry. Four experiments are shown, each including four conditions (G2M: DMSO or 5-ph-IAA; Release G1: DMSO or 5-ph-IAA). The gating strategy is shown.
EdU positive G1 and G2_M are gated.

#### Additional information Figure 3

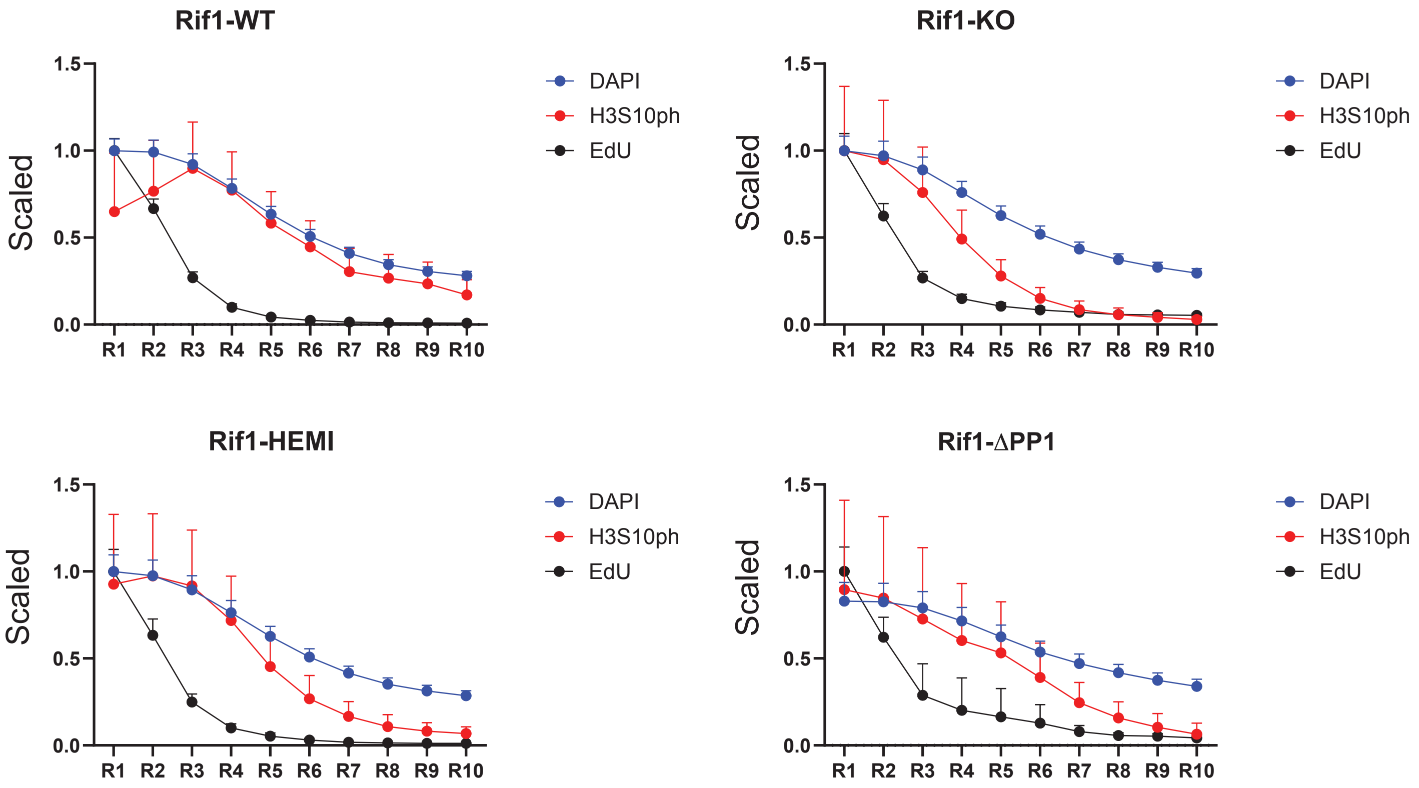

**Additional information Figure3.** Immunofluorescence analysis showing premature, pre-replication phosphorylation of H3S10 in Rif1-KO, Rif1-ΔPP1 and Rif1-HEMI cells. Second, independent experiment for experiment shown in Figure 5F. At least two clones of each genotype were used for the quantification. Rif1-WT, n=3; Rif1-KO, n=2; Rif1-HEMI, n=3; Rif1-ΔPP1, n=4. The plots show the mean and 95% confidence intervals of DAPI, EdU and H3S10ph.

#### Additional information Figure 4

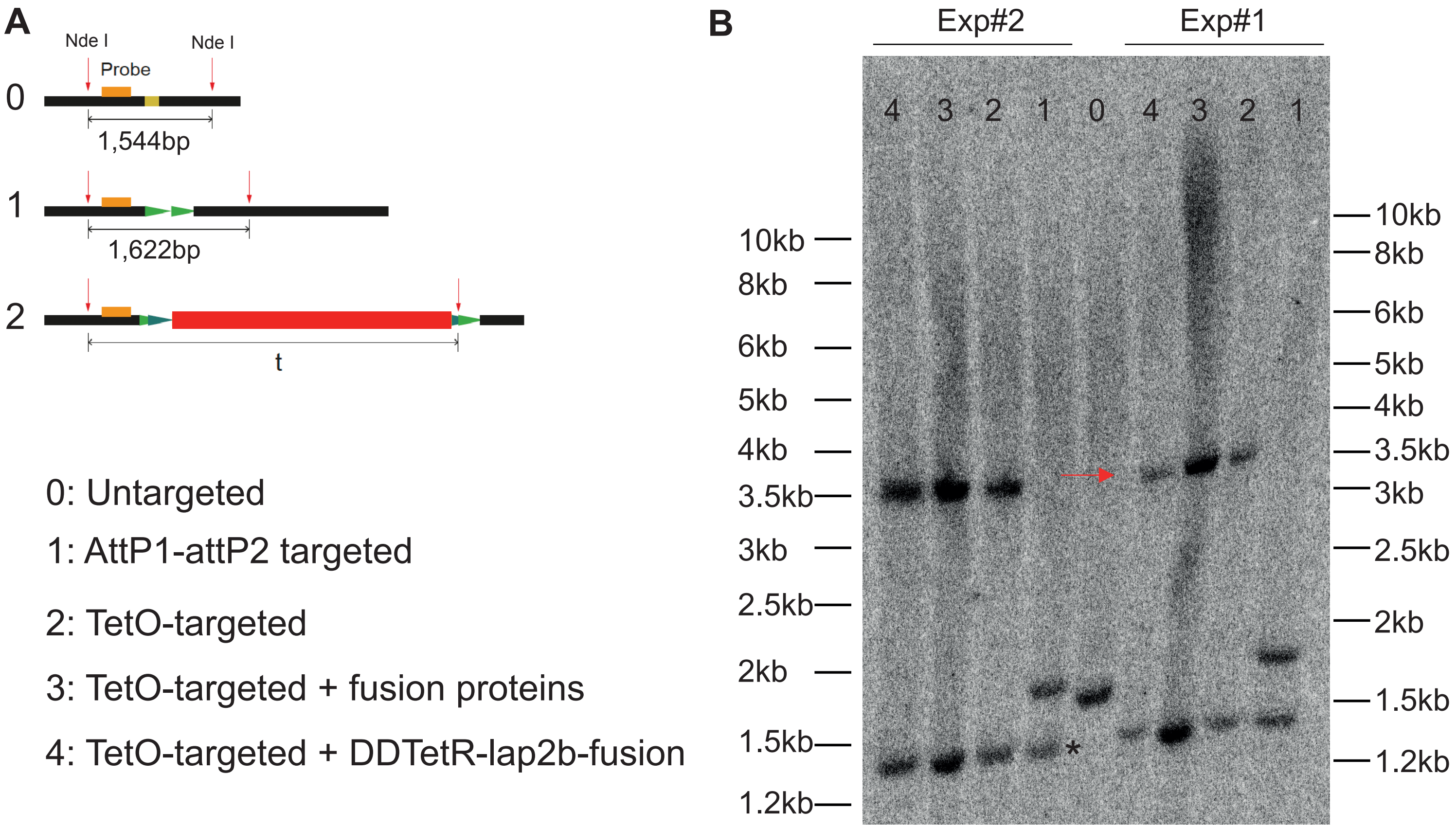

**Additional information Figure 4:** Southern blotting validation of *TetO* arrays targeting in the O6 locus in the O6-Tether cell line. **A**. schematic of the targeting. Black line= genomic DNA; yellow box= targeted site. Green arrowheads=attP1 and P2; red box=*TetO*. **B** O6-Tether cells were generated in a step-wise manner. All intermediate cell lines are shown here. Two batches of samples were collected and processed (exp#1 and 2). Lane 0 shows the original *Rif1*$^{flox/flox}$ *Rosa26*$^{CreERT/+}$ cells (Untargeted) and the expected band at 1,544bp. Lane 1 shows AttP1-AttP2 targeted cells and the expected band at 1,622bp. The blot reveals that only one allele was successfully targeted, while the other allele underwent a small deletion, leading to a shorter band lower than 1.5kb (*). Lane 2 shows the integration of a *TetO* array containing roughly 70 repeats, as calculated by the size of the band. The original array contained 224 repeats and would have given a 12,461bp band. However, the integration of a smaller number of repeats was expected, as deletion within the repeats takes place during the recombination step. In lanes 3 and 4 genomic DNA from cells carrying the *TetO* array and stabling expressing fusion proteins, as shown in Supp. Fig. 4B.The equal length of the 3.5kb fragment in lanes 2, 3 and 4 indicates that the *TetO* arrays is stably maintained.

#### Additional information Figure 5

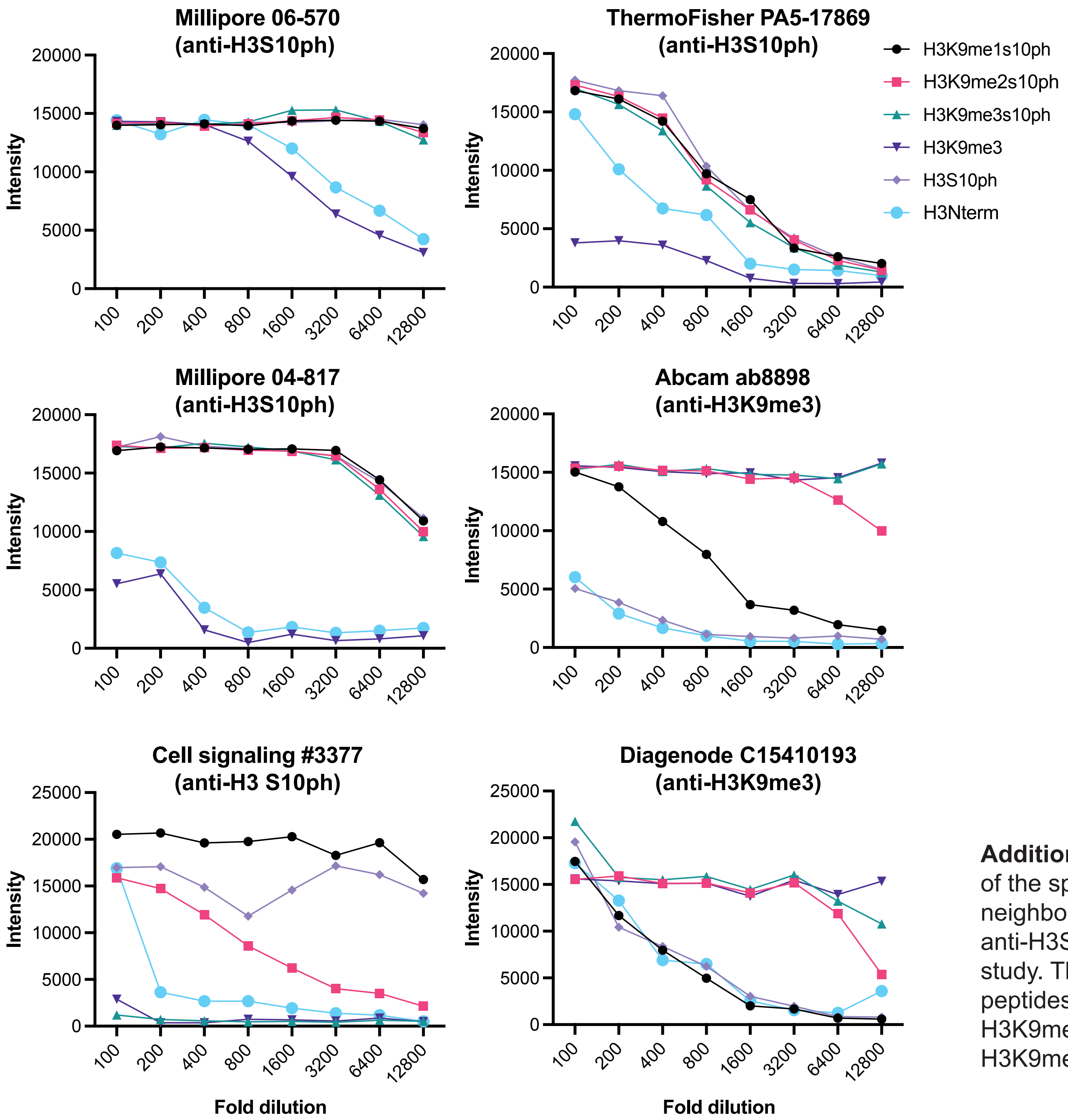

**Additional information Figure 5**: evaluation by ELISA of the specificity and sensitivity to modification of neighbouring residues for a number of different anti-H3S10ph and H3K9me3 antibodies used in this study. The antibodies were tested against different H3 peptides, including H3Nterm(unmodified), H3S10ph, H3K9me3, H3K9me1S10ph, H3K9me2S10ph and H3K9me3S10ph.

### Materials and Methods

#### Cell culture

Mouse embryonic stem cells (ESCs) were cultured at 37°C, 7.5% $CO_2$ on gelatin (Sigma G9391) in KO-DMEM (Gibco 10829-018) containing 12.5% heat-inactivated foetal bovine serum (FBS, PAN Biotech, tested for ESCs), 1% MEM Non-Essential Amino Acid(Gibco, 11140-035), 1% L-Glutamine (Gibco, 25030024), 1% Penicillin/streptomycin (Gibco, 15070063), 0.1mM 2-mercaptoethanol(Gibco, 31350010), 20ng/mL leukaemia inhibitory factor (LIF, EMBL Protein Expression and Purification core facility), 1µM MEK inhibitor PD0325901 and 3µM GSK3 inhibitor CHIR99021 (2i, Cambridge Bioscience) as previously described (Gnan et al., 2021).

To prepare samples for Mass Spectrometry (MS), ESCs were cultured in DMEM (Life Technologies 11995065), containing 12.5% dialysed FCS (Gemini Biosciences FSD500), For SILAC-MS, DMEM was replaced with DMEM for SILAC (Life Technologies A33822), and Arginine (R0: Sigma-Aldrich A8094 or R10: CNLM-539-H1) and Lysine (K0: Sigma-Aldrich L8662 or K8: CK Isotopes CNLM-291-H) isotopes were added to final concentration at 548µM and 648µM, respectively.

#### Tamoxifen-induced deletion of *Rif1*

Multiple clones of ESCs $Rif1^{wt/wt}$ $Rif1^{flox/flox}$, $Rif1^{wt/flox}$, $Rif1^{TgWT/flox}$ *and* $Rif1^{\Delta PP1/flox}$, all containing $Rosa26^{CreERT2}$ were described in Foti et al, 2016 and Gnan et al., 2021 (listed in the table of Key resources, **Table1**). To delete *Rif1*, 2 million of $Rif1^{wt/wt}$ cells or 2.5 million of $Rif1^{flox}$ cells were plated to 10cm dishes in ESC medium with 200 nM 4-hydroxytamoxifen (OHT, Sigma H7904) for 96 hrs, and fresh media was replaced 48 hrs after plating. Deletion was confirmed by genotyping with the primers listed in **Table 2**.

#### mini-AID-mCherry-RIF1

Cas9 (px458, Addgene:48138) and *Rosa26* targeting sgRNA (SP208, a gift from the Pollard Lab (Bressan et al., 2017) expressing vectors were used to target a Flag-OsTIR1F74G (pSB1476) to the *Rosa26* locus in a wild type ESC line (FH6, C57B6J/129Svj background). Clones were validated by genotyping PCR (**Table2**) and by Western blotting. One heterozygous clone, **FH6-T9** (*Rosa26*$^{OsTIR1F74G/WT}$) was chosen, based on the expression level of OsTIR1F74G to target mini-AID-mCherry repair template (**pSB1477**) to *Rif1* N-terminus (pSB1485 and pSB1486). mCherry positive cells were sorted and PCR genotyped (**Table2**). The expression of miniAID-mCherry-Rif1 was validated by Western blotting and immunofluorescence. One homozygous clone (*Rif1*$^{\text{miniAID-mCherry/miniAID-mCherry}}$, *Rosa26*$^{\text{OsTIR1F74G/WT}}$) was used in this study, **RIF1-mAID-11**. For chronic RIF1 degradation, cells were treated with 1μM 5-ph-IAA (diluted from 10mM stock in DMSO, Cambridge Bioscience, HY-134653) or DMSO control for 96 h, and medium was changed every 24 hrs. DMSO or 200nM Hesperadin was added for the last 2 hrs. For acute degradation, 5-ph-IAA was added in the medium at 1μM for 2 hrs or 4 hrs, as well as the DMSO control. The degradation of RIF1 for each experiment was validated by Western blotting.

#### Tethering system

CRISPR-Cas9 system was used to integrate attP1-attP2 landing pad to the genomic site within the FISH probing region O6 (BAC: RP24-318O6, mm10: chr9:116,072,339-116,245,991): O6 targeting sgRNA plasmid (**pSB1210**); repair template **pSB1204**. The *TetO* array (donor plasmid **pSB1196:** attB1 and attB2 at either end of the *TetO* arrays) was inserted in the attP1-attP2 landing pad by co-transfection with the pPGKphiC31obpA-T2A-GFP (**pSB1195**). Clones were screened by genotyping (primers in **Table2**) and validated by Southern blotting (see Additional information Figure 4). The plasmid expressing FRB-Myc-mini-Lap2β-T2A-HA-TetR-FKBPx2-T2A-Puro (**pSB1401**) was randomly integrated, and cells were subcloned to achieve homogenous expression. One *Rif1*$^{flox/flox}$ Rosa26$^{\text{CreERT/+}}$ O6$^{\text{TetO/+}}$ subclone (O6-Tether) was selected to continue the experiments. To induce the tethering, the cells were incubated in medium with or without OHT for 96 hrs to induce *Rif1* deletion. 48 hrs after starting OHT treatment, the medium was replaced with fresh medium containing Rapamycin 10nM (Cambridge Bioscience, SM83-5) or DMSO with OHT.

#### Immunofluorescence

Cells were fixed in 3% w/v paraformaldehyde (PFA) in PBS for 10min at room temperature. If pre-extracted, cells were first incubated in Triton buffer (0.5% Triton X-100, 20mM HEPES, 50mM NaCl, 3mM $MgCl_2$ and 300mM Sucrose) for 2min in the fridge and then fixed in 3% w/v PFA 2% w/v sucrose. Immunofluorescence for H3S10ph was carried out immediately after fixation, to avoid loss of the phosphate group. A number of anti-H3S10ph and anti-H3K9me3 antibodies were screen by ELISA to assess sensitivity to the modification of the nearby residue and the specificity over unmodified H3 or mono and di-methylated 9 (Additional information Figure 5). Only H3S10ph antibodies that did not display interference by the presence of methylation of K9 were selected for use. Millipore 06-570 and CMA313 (Hayashi-Takanaka et al., 2009) were used in this study for Supp.Fig.9D and E. Fixed cells were permeabilised in 0.5% NP40/PBS or Triton buffer (for pre-extracted cells) at room temperature for 10min. Samples were then blocked with PBG buffer (0.2% v/v gelatin and 0.5% w/v BSA in PBS) for 30min at room temperature. Primary antibodies were diluted in PBG and then added to samples for 2hrs incubation at room temperature. Secondary antibodies and DAPI were diluted to 2.5μg/mL in PBG and incubated for 45min at room temperature. Slides were mounted with Vectashield (Vector Laboratories, H-1900) or with Vectashield containing DAPI (Vector Laboratories, H-1200). Images were acquired using a Leica confocal TCS SP5 microscope run by LAS AF Software (Leica) or Zeiss 880 AiryScan run by Zeiss ZEN Black software. Images were cropped and the contrast of images was equally adjusted for all, using ImageJ. Figures were composed in Adobe Illustrator. Detailed information about the antibodies used is listed in the table of Key Resources (**Table1**).

##### EdU staining by Click chemistry coupled with immunofluorescence

Cells were pulse-labelled with 10μM EdU (5-ethynyl-2´-deoxyuridine, Invitrogen, A10044) for the indicated time (standard, 30 min). For pulse-chase experiment, after 30 min EdU pulse, medium was removed and cells were rinsed with pre-warmed PBS twice, followed by incubation in fresh ESC medium to allow chase for indicated time. EdU staining was performed by click chemistry. Click-it reaction cocktail, containing 1x Click-it cell reaction buffer, 4mM $CuSO_4$, 4.8μM Alexa Fluor 647(or 488) azide (Invitrogen) and 1x Click-it cell buffer additive, was added to the coverslips and incubated at room temperature in the dark for 30min. Immunofluorescence was performed as above. Depending on the antibody, the Click-it reaction was performed either before immunofluorescence or after, with a second fixation in PFA between immunofluorescence and Click-it.

#### Analysis of EdU Pulse-chase experiment combined with immunofluorescence for H3S10ph

From each channel the fluorescence background was subtracted by setting an intensity threshold. Next, in the EdU (green) channel the ImageJ findMaxima function was used to mark sites of active DNA replication, adjusting the prominence to make sure that each EdU spot is marked by only 1 maximum. The coordinates for these maxima were stored to perform fluorescence intensity measurements in each channel at the same location. To each maximum a circle was fitted, and the integrated fluorescence intensity per area measured. Subsequently, a series of 9 nested donuts was generated, each having a 3-pixel thickness (106nm) and the integrated intensity per area measured for each. Therefore, 10 data points were obtained for each maximum per channel. Finally, radial intensity profiles were generated by using the donut centre as origin and plotting the average fluorescence intensity as a function of radial distance with each position corresponding to a 106nm increase in radius.

The intensity profile of each fluorescent channel was scaled 0-1 by normalising to the position with the maximum average fluorescence, before plotting profiles of different channels together in GraphPad Prism.

#### FACS sort of live cells

Cells were washed once with FACS buffer (1%FBS/PBS, supplemented with +1% Penicillin/streptomycin), resuspended in FACS buffer at cell density of less than 10million/mL and kept on ice before sorting on a FACSAria (BD). During sorting, gates were drawn based on negative control(s), and cells were sorted into of pre-warmed ESC medium (supplied with 20% FBS, 2i and LIF).

#### FACS analysis of EdU staining

EdU pulsed cells were collected, washed with PBS, 2%FBS/PBS and fixed in 2% PFA/PBS (7.5 million cells/mL) for 10min. at room temperature. Fixed cells were rinsed with 10 volumes of pre-chilled PBS, resuspended in PBS (15 million cells/mL) and kept in the fridge before use. Alternatively, EdU pulsed cells were fixed with cold 70% ethanol and kept in the freezer. For EdU Click-it reaction, fixed cells were permeabilised with 0.05% v/v Triton X-100/PBS for 2min. at room temperature, washed with 10 volumes of cold PBS and pelleted at 400g for 5min. at 4°C. Cells were resuspended in Saponin solution (0.1% w/v saponin, 1% w/v BSA in 1x PBS) at 1.5million/mL for 5min incubation at room temperature. Click-it reaction was performed in 10mM Sodium Ascorbate, 1µM Alexa Fluor 647(or 488) azide and 2mM $CuSO_4$ in PBS for 30min at room temperature in the dark on the nutator. Cells were pelleted at 800g for 5min. and washed with Saponin solution twice. Then cells were resuspended in 2.5µg/mL DAPI/saponin solution at 0.5million cells/mL. Samples were kept in fridge overnight for DAPI staining before analysis using BD LSRFortessa run by BD FACSDiva software. The data was analysed with Flowjo v10 (BD).

##### FACS analysis of H3S10ph immunostaining

After saponin permeabilization supplemented with phosphatase inhibitor (Merck Life Sciences UK, P0044-5ML), anti-H3S10ph antibodies were diluted in 100µL Saponin solution and incubated with 1million cells for 1 hr at room temperature on the nutator. Samples were rinsed three times with 1mL 1% w/v BSA/PBS and spun at 600g for 5 min, at 4°C. Secondary antibodies (table of key resources, **Table1**) were diluted 1:1,000 in saponin solution and added to cells for incubation at room temperature for 1hr in dark. Samples were rinsed three times with 1mL 1% w/v BSA/PBS. Post-staining fixation was performed using 2%PFA/PBS for 10 min, followed by washes with 1% w/v BSA/PBS. Subsequently, cells were resuspended in 400µL of saponin solution with DAPI 2.5µg/mL. Samples were kept in the fridge overnight for DAPI staining and analysed with BD LSRFortessa run by BD FACSDiva software, after filtration with 35µm nylon mesh to 5mL polypropylene round-bottom tubes. The analysis was performed with Flowjo v10 (BD).

Flow cytometry analysis for SuppFig9 hesperadin treatment: the mean intensity of DAPI, H3S10ph of gated intact cells and EdU negative G1 cells were exported from FlowJo. Then for each sample, the DAPI intensity (intact cells) was normalised to the mean DAPI intensity of G1 cells. The density plots were made by ggplot in R.

##### FACS sorting of fixed cells, followed by cytospin and H3S10ph immunofluorescence.

Cells were stained for EdU as described above. Samples were filtered with 35µm nylon mesh and sorted into 4 fractions based on DAPI staining. 50,000 sorted cells were cytospun with Shandon cyto-funnels (Thermo Scientific 12016679) at 1,800rpm for 5min. onto polysine slides (VWR 631-1349), and then fixed again in 4% w/v PFA/PBS for 5min. After wash with PBS, the samples were permeabilised again with 0.5% v/v NP40/PBS for 10 min. Following PBS washes, Click-it reaction and immunofluorescence were performed as described.

##### Analysis of H3S10ph immunofluorescence in FACS -sorted and cytospun cells

Individual nuclei were segmented using ImageJ. Then G1 nuclei were identified with negative EdU signal in sorted P1, while early S-phase nuclei were identified with positive EdU in sorted P2. Nuclei with H3S10ph signal at chromocenters were manually scored (at least 50 nuclei were included each sample). The plots were made in GraphPad Prism.

##### Western blot

Cells were lysed in 2x Leammli buffer (90mM Tris-HCl pH6.8, 1.85% SDS, 18.5% glycerol, 0.5% Bromophenol blue 2.1% v/v b-mercaptoethanol) at 10,000 cells/μL. After boiling for 10min, the samples were sheared through a 0.22μm syringe 5 times. SDS-PAGE gels were transferred to 0.45μm Nitrocellulose membrane (GE Healthcare Life Science). Membranes were washed with 0.05% Tween 20/PBS (PBST) or 0.1%Tween 20/TBS(TBST) buffer and blocked in 5% milk/PBST or 5% milk/TBST. Primary antibodies were diluted in the blocking buffer. Secondary antibodies (IRDye 680RD or IRDye 800CW, LI-COR, Lincoln, NE, USA) were diluted 1:15,000 in 2.5% milk for 45min. in the dark at room temperature. Membranes were imaged on a LI-COR Odyssey Imager (LI-COR, Lincoln, NE, USA).

##### Immunofluorescence combined with 3D-FISH

ESCs grown on slides were fixed in 4% w/v PFA/PBS for 10min, permeabilised in 0.5% Trition X-100/PBS at room temperature for 10min, and de-hydrated in 70% Ethanol (5min, three times). Immunofluorescence was carried as described above, with a modification of the blocking buffer (1% w/v BSA/0.2% Tween 20/PBS). After immunofluorescence cells were PFA-fixed again. The slides were rinsed with 2x SSC (Life Technologies AM9765) for 5min and then treated with 0.25mg/mL RNase A (Sigma, R5250) in 2x SSC for 1hr at 37°C. After 2x SSC washes, slides were treated with 0.6% HCl and 0.06% v/v Tween 20 for 10min, at room temperature, rinsed with 0.2% Tween 20/2x SSC and 2x SSC at room temperature for 5min and dehydrated sequentially in 70%, 80%, 90%, 100% ethanol. Air dried samples were denatured in 50% formamide (Sigma-Aldrich, 47671) in 2xSSC at 85°C (pH7.5) for 30min. Following denaturation, the slides were dehydrated in pre-chilled (-20°C) ethanol, 70%, 80%, 90%, 100%, for 5min each. Digoxigenin (DIG)-labelled probes (BACs in Table 1)(prepared according to manufacturer's instruction, Roche, 11745816910) mixed with Cot1 DNA (Life Technologies,18440016) and 1µg of Salmon Sperm DNA (Invitrogen, AM9680) were precipitated and then denatured in 100% formamide (Sigma, F9037a) at 80°C for 7min. and cooled to 37°C by 0.1°C/s. Hybridisation buffer (20% dextran sulphate, 0.4% BSA, 4xSSC) was added to probe in a 1:1 v/v ratio and applied to air-dried de-hydrated cell samples. Following further denaturation for 2min at 85°C, samples were incubated in a humid chamber at 37°C overnight. Washes in 50% formamide in 2xSSC (pH7.5) (5min, three times, the first at 45°C, the next two at room temperature) and 2xSSC (5min, three times). Blocking was carried out for 30min, with 3% w/v BSA/0.2% Tween 20/PBS at room temperature. Then the slides were incubated with anti-DIG-488 (Jackson Immunoresearch 200-542-156) 1:500 in 1% BSA w/v /0.2% Tween 20/PBS, at 37°C for 1hr. After 3 washes of 0.2% Tween 20/PBS and 3 washes of PBS, the slides were mounted with Vectashield (Vector Laboratories, H-1200). Images were acquired using a Leica confocal TCS SP5 microscope run by LAS AF Software (Leica), with z-stacks 0.3µm.

##### Immunofluorescence-FISH lamina association frequency analysis

Lamina-association was manually classified while navigating z-stacks. When at least one allele was located at the nuclear periphery, overlapping with LaminB1 signal, the nucleus was classified in the lamina-associated category. Overlap with Lamin B1 was defined as the shortest distance between the centre of FISH signal and LaminB1 signal <0.45µm=9pixels. By going through the Z-stacks, the stack where the FISH signal appeared brightest was chosen for classification. More than 80 nuclei are included for classification per sample for each experiment. The classification was then combined from different experiments. The plots were made in GraphPad Prism. The statistical analysis was performed with two-tailed Welch's *t*-test.

#### Dual-colour FISH

Cells were resuspended in 5mL of hypotonic buffer (75mM KCl), added dropwise while gently vortexing, incubated at 37°C for 10min and pelleted at 1,200rpm (262g) for 5min. at room temperature. The pellets were fixed in fixative (3:1 methanol: glacial acetic acid) 3 times, spinning at 1,200rpm(262g) for 5min. Cells were added to slides washed in ethanol and HCl, and airy dried. DIG-BAC and Biotin-TetO probes were prepared following manufacture's instruction (DIG or Biotin NICK TRANS MIX, Roche). The two probes were mixed at 1:1 and precipitated. The probe-mix was resuspended in hybridisation buffer (50% Formamide, 2xSSC, 1% Tween 20, 8% Dextran sulphate). The slides were incubated in 2xSSC with 100μg/mL RNase A at 37°C for 1hr, washed with 2xSSC, dehydrated in 70%, 90% and 100% ethanol in series, 2min each. Air dried slides were denatured at 70°C for 5min then moved to 70%Formamide/2xSSC (pH7.5) for 2min. Sequential washes were performed with pre-chilled ethanol series, 70%, 90% and 100%, 2min. each. The probe-mix was denatured at 70°C for 5min. and added to each slide dried and pre-warmed on heat block at 37°C. The hybridisation was incubated in humid chamber at 37°C, overnight. Slides were subsequently washed in 2xSSC at 45°C (3min, 4 times), followed by washes in 0.1%SSC at 60°C (3min, 4 times), rinsed in 0.1%Tween 20/4xSSC at 37°C (3min, 3 times). Slide were incubated in blocking solution (5% milk in 4xSSC, prewarmed in 37°C water bath for 10min) for 10min at 37°C. Anti Digoxigenin-Rhodamine (Roche, 11207750910) was diluted in blocking buffer 1:200, as first layer; Avidin-FITC (Vector Labs, A-2011) 1:200 and Texas red anti Sheep (Vector Labs, TI-6000) 1:100 were the second layer; Biotin anti Avidin (Vector Labs, BA-0300) 1:200 was the third layer. Avidin-FITC (Vector Labs, A-2011) 1:200 was the last layer. For each layer, the incubation was at 37°C for 45min. Between layers, the slides were rinsed with 0.1% Tween 20/4xSSC (3min., for 4 times) and finally mounted with Vectashield with DAPI.

#### Southern blotting

2 million cells were lysed in 500μL of digestion buffer (50mM TrisHCl pH8, 100mM EDTA, 100mM NaCl and 1% SDS) with 800μg/mL proteinase K (Sigma-Aldrich, P6556) and incubated at 54°C overnight. DNA extracted by phenol-chloroform was digested with the selected restriction enzymes at 37°C overnight. 1.5-2.5μg of digested DNA was run on a 0.7% agarose gel. The gel was blotted onto Hybond N+ membrane (GE Healthcare Life

Sciences, RPN 203 S). The membrane was brought back to pH 7- 7.4 by rinsing in SCC2X and blocked in pre-hybridisation buffer (0.5M $Na_2HPO_4$, 1mM EDTA, 5% w/v SDS, 3% w/v BSA in $ddH_2O$) at 65°C for about 2hrs. Probes were prepared by PCR using specific BACs (table of key resources, **Table1**) as templates with the primers listed in **Table2**. α-P32-dATP (Perkin Elmer, BLU012Z) was incorporated into the denatured probe by Klenow polymerase, in a reaction containing dCTP, dGTP, dTTP and α-P32-dATP (Random primed DNA labelling kit, Roche-11004760001). Subsequently, the radioactive isotope-labelled probe was denatured at 95°C for 3min. and used for hybridisation at 65°C. Washing buffer (40mM $Na_2HPO_4$, 1mM EDTA, 5% w/v SDS in $ddH_2O$) was used at 65°C for 20min, three times. Membranes were exposed overnight to a cyclone screen, then scanned in Typhoon FLA 7000 scanner. The contrast of images was adjusted in ImageJ. Images were cropped in ImageJ before figure assembly in Adobe Illustrator.

##### H3K9me3 ChIP-seq

Cells in OHT for 4 days were enriched at G1/S by incubation in 2mM thymidine (Sigma-Aldrich, T9250) for 15hrs before collection. This eliminated the potential variability of H3K9me3 levels due to the accumulation of Rif1-KO, Rif1-HEMI and Rif1-ΔPP1 cells in G2 (Gnan et al., 2021). Pelleted cells were crosslinked in 1% Formaldehyde (Sigma-Aldrich 252549) in crosslinking solution (50mM HEPES pH7.9, 150mM NaCl, 1mM EDTA, 0.5mM EGTA in PBS) for 10min at room temperature, followed by quenching with 0.125M glycine for5 min. Cells were pelleted at 1,400rpm (359g) for 5min, at 4°C , washed with pre-chilled DPBS supplemented with 0.3mM PMSF. Cells lysed overnight in lysis buffer (1% SDS, 10mM EDTA, 50 mM Tris-HCl pH 8, supplemented with 1x Complete EDTA-free Protease inhibitor (Sigma, 5056489001)) were then sonicated in a Bioruptor sonicator at 4°C to the desired range 200-300bp. Fragmented chromatin was pre-cleared by centrifugation at 400g for 20min at 4°C. Qubit dsDNA HS assay kit (Q32854) was used to quantify the chromatin. For each immunoprecipitation, 10μg of chromatin were diluted in 10 volumes of dilution buffer (1% Triton x100, 2mM EDTA, 167mM NaCl, 20mM Tris-HCl pH8.1, supplement with 1x protease inhibitor), and incubated with 1.6μg of antibody (H3K9me3, ab8898 or rat immunoglobulin G (IgG), sc-2026, Santa Cruz) overnight rotating at 4°C. 1% of the chromatin was taken as input control. The following day, 50μL of protein G Dynabeads (Invitrogen, 10004D) were added to each ChIP sample for 2hrs incubation at 4°C in rotation. The beads were washed in low salt buffer (0.1% SDS, 1% Triton, 2mM EDTA, 150mM NaCl, 20mM Tris-HCl pH8.1) three times, followed by one wash in high salt buffer (0.1% SDS, 1% Triton, 2 mM EDTA, 500mM NaCl, 20mM Tris-HCl pH8.1), one wash in LiCl buffer (0.25M LiCl, 0.5% NP-40, 0.5% sodium deoxycholate, 1mM EDTA,10mM Tris-HCl pH 8.1) and one wash in TE (10mM Tris, 1mM EDTA, pH8). All wash buffers were supplemented with 1x protease inhibitors and every wash was 5min in rotation at 4°C. After the final wash in TE buffer without protease inhibitor, chromatin was eluted by incubating beads with Elution buffer (1% SDS, 100 mM NaHCO3) for 1hr rotating at room temperature. Beads collected on the magnet and then the supernatant was transferred and incubated with RNase A (to final at concentration of 187.5μg/mL, Sigma R5250) for at least 1hr at 37°C, followed by addition of proteinase K (at final at 225μg/mL, Sigma-Aldrich P6556) and overnight incubation at 60°C for reverse-crosslinking. Input samples were processed in parallel for RNase and proteinase K treatment.  Input and IP samples were cleaned up using Zymo ChIP DNA clean and concentrator kit (Zymo research, D5205). Libraries were prepared using NEBNext® Ultra™ II DNA Library Prep Kit (NEB, E7103L) following the manufacture's instruction. The libraries were amplified with PCR using index primers, for 4 cycles, followed by purification with SPRIselect® Reagent Kit (Beckman Coulter, Inc. #B23317). (GEO: GSE299608, see also table of Key resources, **Table1**)

#### ChIP-Seq analysis

Data processing of fastq files was performed using a reproducible ChIP-seq workflow implemented with Snakemake (v6.15.1). ChIP-seq libraries were sequenced using paired-end Illumina chemistry with a read length of 2× 50b. Adapters were trimmed using Cutadapt v1.18(Martin, 2011) with --minimum-length 20 and --nextseq-trim=20. Quality of raw and trimmed reads was assessed using FastQC v0.11.9 (http://www.bioinformatics.babraham.ac.uk/projects/fastqc/) and summary reports were aggregated using MultiQC v1.26(Ewels et al., 2016). Trimmed reads were aligned to the mouse mm10 reference genome using BWA-MEM v0.7.16(Li and Durbin, 2009) with the -M flag to mark shorter split hits as secondary. The resulting SAM files were filtered, sorted and indexed using Samtools v1.17￼(!!! INVALID CITATION !!! ￼, and alignment statistics were collected via samtools)flagstat. Alignments were subjected to the following filtering steps to retain a single alignment for each mapped read: Removal of secondary (0x100), supplementary (0x800), and unmapped (0x4) reads (-F 2304), retention of properly paired reads (-f 3). Duplicate reads were removed using Picard MarkDuplicates v3.3.0(https://broadinstitute.github.io/picard/) and reads aligning to a mm10 blacklist (https://www.encodeproject.org/files/ENCFF547MET/) were removed with bedtools intersect v2.3.0 (Quinlan and Hall, 2010).

Signal tracks were generated using deepTools v3.5.0(Ramirez et al., 2016). bamCoverage was used to compute normalised coverage in bins of 1 bp using the BPM (bins per million reads) method. bamCompare was used to compute log2 enrichment profiles for IP over input control, using exact scaling and BPM normalisation.

##### ChIP-seq peak calling

Peaks of H3K9me3 were called with MACS2 callpeak v2.2.7.1(Zhang et al., 2008) using the corresponding Input sample as control with uniquely aligned reads only. Differential H3K9me3 peaks between WT and KO were called using DiffBind 3.8.4 with an adjusted p-value threshold of 0.05 and a log fold change > 0.75.

##### Genomic overlap analysis

Overlaps between genomic features, repeat regions, ChIP-seq peaks and domains were calculated using the R programming language. The RegioneR package 1.3.0 (Gel et al., 2016) was used to perform pemutation tests to determine whether the cumulative coverage of overlapping regions was statistically significant. Permutation tests were run 1000 times, or 100 times, depending on the number of regions involved.

##### Read counts over repeats

Read counts across different repeat annotations were quantified with featureCounts v2.0.3(Liao et al., 2014) using the parameters -p -M to include multi-map reads. RepeatMasker annotations for mm10 were downloaded from the UCSC database (http://hgdownload.soe.ucsc.edu/goldenPath/mm10/database/rmsk.txt.gz) and converted to gff with the repeat name as gene_id to cumulatively count reads aligned to each repeat type.

##### Data visualisation and plotting

Plots of ChIP-seq read counts and overlaps were generated in R with the tidyverse packages(Wickham et al., 2019). Snapshots of ChIP-seq read profiles are taken from the Integrative Genomics Viewer (IGV)(Robinson et al., 2011).

#### Hi-C analysis

TADs Clustering: The list of TADs was generated using the HiC data from Gnan et al., 2021. TADs were clustered by intensities of LaminB1 and RIF1, individually, by k-means clustering using deepTools. The resulting 3 groups defined by LaminB1 intensity and 2 groups by RIF1 intensity were joint to create 6 clusters in total. The heatmaps were made by deepTools (Galaxy). To make box plots, HiC Eigenvector, H3K9me3 ChIP, replication timing profiles were first mapped to TADs by bedtools, calculating the mean intensity. Plots were made using seaborn in Jupyter notebook.

#### qPCR

ChIP DNA was diluted to same concentration in each experiment. qPCR was performed by using LightCycler® 480 SYBR Green I Master (Roche 04887352001), following manufacturer's instructions on a LightCycler® 96 Instrument (Roche).

#### Repli-seq

Rapamycin-induced tethering in ESC O6-Tether, as described above. Before collection, BrdU was added to the culture medium to a final concentration of 200μM for 1hr. Cells were washed in 1% v/v FBS/DPBS and resuspended at 15-20million cells/2.5mL 1% v/v FBS/DPBS. 7.5mL of ice-cold 100% ethanol was added drop-wisely with gentle vortexing to fix cells. Following FACS sorting in four fractions, NGS libraries preparation and BrdU immunoprecipitation followed the procedures described in (Zhao et al., 2020, Marchal et al., 2018). The first two fractions were pooled as the early S-phase fraction, while the latter two fractions were pooled for the late S-phase fraction.

#### Repli-seq analysis

Repli-seq data were analysed as described in Marchal et al., 2018. The reference sequence for Chr9 was rebuilt to include the *TetO* integration. The replication timing profiles were visualied using IGV.

#### ELISA

96-well plates (Pierce Streptavidin High Binding capacity coated 96-well plate, #15503) were coated with 100μL of biotinylated peptides per well at the concentration of 1μg/mL, diluted in coating buffer (50mM sodium carbonate/bicarbonate, pH 9.4). (Peptides from Epicypher Epi H3 – cat # 12-0001, H3K9me3 cat # 12-0012, H3K9me3S10ph -cat # 12-0054, H3K9me1S10ph- cat # 12-0093, H3K9me2S10ph- cat # 120155,H3S10ph – cat # 12-0041). The incubation was carried out at 4˚C overnight. The plates were washed three times with wash buffer (TBS-0.05% Tween 20) on a shaking platform to remove unbound peptides. 300uL of blocking buffer was added per well and the plates were incubated at room temperature for 1hr. Subsequently, antibodies (**Table1**) were diluted in series and 100μL of each dilution was added to wells. After incubation at 37˚C for 1hr, the plates were washed three times with wash buffer. Secondary antibodies were diluted and added to plates for 1hr incubation at room temperature. After three washes, 100μL of QuantaRed Working solution was added to each well and incubated for 15min, at room temperature. Stop peroxidase activity by adding 10μL of QuantaRed Stop solution and shake plates for 10-15s. Measure relative fluorescence units (RFU) of each well. The excitation and emission maxima of QuantaRed Substrate are 570nm and 585nm, respectively, recorded on a colorimetric plate reader Spectra max M5 from Molecular Devices.

#### iPOND-SILAC

RIF1-mAID-11 cells were synchronised in 2mM thymidine in light SILAC medium (R0K0) overnight (around 15hrs), then released to heavy SILAC medium (R10K8) supplemented with Nocodazole (50ng/mL) for 8hrs. EdU (10μM) was added at multiple time points (every 50min.) for 15min, between 3 and 8hrs after release from thymidine to label chromatin replicated in mid-late S phase. When cells were in G2/M (8hrs after thymidine release), DMSO or 5-ph-IAA (1μM) were added for 2hrs to degrade RIF1, followed by 2hrs release from Nocodazole. Cells were crosslinked with 1% Formaldehyde (Sigma-Aldrich 252549) for 15min. and quenched in 0.125M Glycine for 5min. Crosslinked cells were pelleted by centrifugation at 1000rpm (182g) at 4°C. After wash with PBS, cells were pelleted again and were snap-frozen in liquid nitrogen. iPOND was performed as described previously (Sirbu et al, 2012). Cells were permeabilized in 0.25% Triton X-100 in PBS for 30min at room temperature and washed with 0.5% BSA/PBS. To conjugate biotin to EdU-labelled DNA, click chemistry with Biotin-azide was performed for 1.5hrs in the dark (10µM Biotin-azide (Thermo Fisher, B10184), 10mM Sodium ascorbate, 2mM $CuSO_4$ in 1×PBS). Cells were lysed (1% SDS in 50mM Tris-HCl, pH8.0, protease inhibitor), sonicated (Bioruptor, Diagenode). Then the lysate was diluted 1:1 (v/v) with cold PBS containing protease inhibitor, and the biotin-conjugated DNA-protein complexes were captured by streptavidin beads (Thermo Fisher, 65002). The captured complexes were washed with lysis buffer and 1M NaCl, followed by three washes with 50mM Tris-HCl, pH 8. The protein-conjugated beats were dried using a vacuum centrifuge and further processed by EpiQMax.

Beads were processed according to a SP3 protocol as described previously (Hughes et al., 2019). However, proprietary steps developed by MOLEQLAR Analytics GmbH have been added in order to adjust the protocol for histone specific aspects. The protocol started with protein propionylation. Upon 3hrs digestion at 37°C and 1,000rpm in a table-top thermomixer, samples were acidified by adding 5µL of 5% TFA (trifluoroacetic acid) and quickly vortexed. Beads were immobilized on a magnetic rack, and peptides were recovered by transferring the supernatant to new tubes. Samples were dried down using a vacuum concentrator and reconstituted by adding 20µL of 0.1% formic acid to reach a peptide concentration of approximately 0.2µg/µL. MS injection-ready samples were stored at -20°C.

##### LC-MS analysis of histone modifications

Approximately 200ng of peptides from each sample were separated on a C18 column (Aurora Elite TS, 15cm x 75µm ID, 1.7µm, IonOpticks) with a gradient from 5% B to 25% B (solvent A 0.1% FA in water, solvent B 100% ACN, 0.1% FA) over 35min. at a flow rate of 300nl/min (Vanquish Neo UHPLC-Systems, Thermo - Fisher, San Jose, CA) and directly sprayed into a Exploris 240 mass spectrometer (Thermo-Fisher Scientific). The mass spectrometer was operated in full-scan mode to identify and quantify specific fragment ions of N-terminal peptides of human histone 3.1 and histone 4 proteins. Survey full scan MS spectra (from m/z 250–1600) were acquired with resolution 60,000 at m/z 400 (AGC target of 3x106). Typical mass spectrometric conditions were: spray voltage, 1.9kV; no sheath and auxiliary gas flow; heated capillary temperature, 300°C.

##### iPOND-MS

For iPOND-MS cells were synchronised and released the same as iPOND-SILAC experiment, but only light medium was used. iPOND was performed as described above, except that captured complexes were washed extensively using lysis buffer and 1M NaCl. The proteins were eluted under reducing conditions by boiling in 2× LSB sample buffer for 30min.

##### Tandem mass tag (TMT)-based MS for quantitative proteomics analysis

iPOND samples were prepared using SP3 protocol for TMT labelling as described in (Hughes et al., 2019). In brief, proteins were eluted in 2xLSB buffer. After the samples were denatured at 95℃ for 10 min., they were alkylated with 400mM IAA (final concentration) in dark at room temperature for 30 min. Protein concentration was measured using EZQ™ Protein Quantitation Kit by following the manual. Then, protein samples were mixed and cleaned with SP3 beads (1: 10, protein:beads), and digested with lysC/trypsin mixture (1: 50, enzyme:protein). 100μL 100mM TEAB buffer (pH8.0). was used to elute the peptides from SP3 beads.

All eluted peptides from each sample were labelled with TMT11plex Isobaric Mass Tag Labelling Kit (Thermo Fisher Scientific) by following it's manual. A pool of peptides from all Input samples were created and labelled with one TMT channel as the reference channel for normalisation. TMT labelled peptide samples were further mixed and fractionated using offline high-pH reverse-phase (RP) chromatography, as previously described (Brenes et al., 2019). 24 fractions were subsequently collected and dried in SpeedVac. Peptides were re-dissolved in 5% formic acid.

TMT fractions samples were analysed by using an Orbitrap Fusion Tribrid mass spectrometer (Thermo Fisher Scientific), equipped with a Dionex ultra-high-pressure liquid-chromatography system (RSLCnano). RPLC was performed using a Dionex

RSLCnano HPLC (Thermo Fisher Scientific). Peptides were injected onto a 75μm×2cm PepMap-C18 pre-column and resolved on a 75μm×50cm RP- C18 EASY-Spray temperature-controlled integrated column-emitter (Thermo Fisher Scientific), using a 120-min multistep gradient from 10% B to 44% B with a constant flow rate of 300nl/min. The mobile phases were: H2O incorporating 0.1% FA (solvent A) and 80% ACN incorporating 0.1% FA (solvent B). The data were acquired under the control of Xcalibur software in a data-dependent acquisition mode (DDA) using top speed and 3 s duration per cycle. The survey scan was acquired in the orbitrap covering the m/z range from 375 to 1,600Da with a mass resolution of 120,000 and a stand automatic gain control (AGC) target. The most intense ions were selected for fragmentation using CID in the ion trap with 35% CID collision energy and an isolation window of 0.7Da. The AGC target was set to standard with automatic maximum injection time and a dynamic exclusion of 60 s. The MS2 scan was acquired in the ion trap in Turbo mode. SPS-MS3 mode was applied for more accurate TMT quantifications. 10 fragment ions were co-isolated using synchronous precursor selection with a window of 2Da and further fragmented using HCD collision energy of 65%. The fragments were then analysed in the orbitrap with a resolution of 50,000. The AGC target was set to standard, and the maximum injection time was set to 105 ms.

#### MS analysis

Quantification of histone modifications (EpiQMAX*)*

Data analysis was performed with Skyline (version 23.1.0.455)(MacLean et al., 2010) by using doubly and triply charged peptide masses for extracted ion chromatograms (XICs). Peaks were selected manually. Heavy arginine-labelled spiketides (13C6; 15N4) were used to confirm the correct retention times and for signal normalization purposes, because all heavy standards were incorporated across all samples at the same concentration. The SILAC peptides (*i.e.* heavy arginine-labelled peptides (R6 Arginine 13C6)) were selected to assess the newly heavy arginine incorporation into synthetized histones proteins. Integrated peak values (Total Area MS1) were used for further calculations. Endogenous PTM signals were normalized according to the variation of the signals of the spiked-in heavy standards and using median normalization. The percentage of each modification within the same peptide is derived from the ratio of this structural modified peptide to the sum of all isotopically similar peptides. Therefore, the Total Area MS1 value was used to calculate the relative abundance of an observed modified peptide as percentage of the overall peptide. The unmodified peptide of histone 3.1 (aa 41–49) was used as indicator for total histone 3.1. Coeluting isobaric modifications were quantified using three unique MS2 fragment ions. Averaged integrals of these ions were used to calculate their respective contribution to the isobaric MS1 peak (e.g., H3K36me3 and H3K27me2K36me1). Plots were generated using GraphPad Prism.

For each condition, the proportion of new H3 histones was calculated per replica, as Alabert et al., 2015.

$$\frac{\textit{sum heavy H3 peptides intens}}{\textit{sum heavy H3 peptides intens} + \textit{sum light H3 peptides intens}} \text{x}100\%$$

LC-MS/MS data analysis

The TMT-MS data were analysed using **MaxQuant (v 1.6.7.0, 10.1038/nprot.2016.136)** and searched against Mus musculus database from Uniport (Swiss-Port and TrEMBL, downloaded in February 2024). The FDR threshold was set to 1% for each of the respective Peptide Spectrum Match (PSM) and Protein levels. The data was searched with the following parameters: quantification type was set to Reporter ion MS3 with TMT 11plex, stable modification of carbamidomethyl (C), variable modifications, oxidation (M) and acetylation (protein N terminus), with maximum of 2 missed tryptic cleavages threshold. The detailed parameter file was uploaded together with the raw MS data.

MS data processing in Perseus/ R-Studio

In Perseus(Tyanova et al., 2016) **(version 2.1.1.0),** the raw dataset was filtered to exclude proteins identified by site, reverse sequences and potential contaminants. For subsequent analyses, the intensities were first transformed to log2 scale and then normalised to the median. In addition, proteins with fewer than two unique peptides were removed. The PCA plot unveiled pronounced batch effects, predominantly associated with distinct mass spectrometer runs. To mitigate these effects, we used HarmonizR(Voss et al., 2022) as plugin within Perseus, utilising the combat method. The subsequent PCA analysis as well as the calculation of the Coefficient of Variation (CV) verified the substantial attenuation of these batch effects. The protein list was then further filtered to retain proteins detected in at least three out of four time points, resulting in a final list of 2849 proteins. Missing values were imputed using K-nearest neighbors (KNN) imputation, which was performed in Perseus for each time point separately (K=3). Differential protein expression analysis was performed using the Limma package (Ritchie et al., 2015) in R, with *p*-values adjusted by the Benjamini-Hochberg (Benjamini and Hochberg, 1995) procedure to control the false discovery rate. Proteins with adjusted *p*-value < 0.05 are considered significant**.** Finally, volcano plots were generated to visualize the results of the differential analysis using R package EnhancedVolcano.

| Reagent or resource | Source | Identifier |
|---|---|---|
| Antibodies | | |
| anti-H3K9me3 | Abcam | Cat#ab8898, RRID:AB_306848 |
| anti-H3S10ph | Millipore | Cat# 06-570, RRID:AB_310177 |
| anti-H3S10ph | Hayashi-Takanaka, Yoko et al., 2009 | CMA311, CMA313 |
| anti-RIF1 | Buonomo et al., 2009 | 1240 |
| anti-Tubulin | Sigma-Aldrich | Cat# T4026, RRID:AB_477577 |
| anti-Suv39h1 | Cell Signaling Technology | Cat# 8729, RRID:AB_10829612 |
| anti-AIM1 | BD Biosciences | Cat# 611083, RRID:AB_398396 |
| anti-SMC1 | Bethyl | Cat# A300-055A, RRID:AB_2192467 |
| anti-MCM3 | Santa Cruz Biotechnology | Cat# sc-9850, RRID:AB_2142269 |
| anti-LaminB1 | Abcam | Cat# ab16048, RRID:AB_443298 |
| anti-Digoxin | Jackson ImmunoResearch Labs | Cat# 200-542-156, RRID:AB_2339039 |
| anti-H4 | Abcam | Cat# ab7311, RRID:AB_305837 |
| anti-SetDB1 | Proteintech | Cat# 11231-1-AP, RRID:AB_2186069 |
| anti-G9a | Cell Signaling Technology | Cat# 3306, RRID:AB_2097647 |
| anti-GLP | R and D Systems | Cat# PP-B0422-00, RRID:AB_2097494 |
| anti-H3 | Cell Signaling Technology | Cat# 3638, RRID:AB_1642229 |
| anti-HA | Covance | Cat# MMS-101P, RRID:AB_2314672 |
| Anti-Myc | Millipore | Cat# 05-419, RRID:AB_309725 |
| anti-FLAG | Agilent | Cat# 200472, RRID:AB_10596649 |
| anti-PP1a | Abcam | Cat# ab52619, RRID:AB_2170391 |
| anti-Digoxigenin Rhodamine | Roche | Cat# 11207750910, RRID:AB_514501 |
| anti-BrdU | BD Biosciences | BD#555627 |
| Rabbit IgG antibody | Epicypher | Cat# 13-0042, RRID:AB_2923178 |
| Rat IgG antibody | Santa Cruz Biotechnology | Cat# sc-2026, RRID:AB_737202 |
| Donkey anti-Mouse IgG (H+L), Alexa Fluor 488 | Thermo Fisher Scientific | Cat# A-21202, RRID:AB_141607 |

| Reagent or resource | Source | Identifier |
|---|---|---|
| Donkey anti-Mouse IgG (H+L), Alexa Fluor 568 | Thermo Fisher Scientific | Cat# A10037, RRID:AB_11180865 |
| Donkey anti-Rabbit IgG (H+L), Alexa Fluor 488 | Thermo Fisher Scientific | Cat# A-21206, RRID:AB_2535792 |
| Donkey anti-Rabbit IgG (H+L), Alexa Fluor 568 | Thermo Fisher Scientific | Cat# A10042, RRID:AB_2534017 |
| Donkey anti-Goat IgG (H+L), Alexa Fluor 568 | Thermo Fisher Scientific | Cat# A-11057, RRID:AB_2534104 |
| Donkey anti-Goat IgG (H+L), Alexa Fluor 647 | Thermo Fisher Scientific | Cat# A-21447, RRID:AB_2535864 |
| anti-Mouse IgG | Sigma-Aldrich | Cat# M7023, RRID:AB_260634 |
| Texas Red anti-sheep | Vector Laboratories | Cat# TI-6000, RRID:AB_2336219 |
| Biotin anti-Avidin | Vector Laboratories | Cat# BA-0300, RRID:AB_2336108 |
| Avidin-FITC | Vector Laboratories | Cat# A-2011, RRID:AB_2336456 |
| IRDye 680RD Donkey anti Rabbit | LI-COR Biosciences | Cat# 925-68073, RRID:AB_2716687 |
| IRDye 680RD Donkey anti Mouse | LI-COR Biosciences | Cat# 925-68072, RRID:AB_2814912 |
| IRDye 800CW Goat anti Rat | LI-COR Biosciences | Cat# 926-32219, RRID:AB_1850025 |
| IRDye 800CW Donkey anti Mouse | LI-COR Biosciences | Cat# 926-32212, RRID:AB_621847 |
| IRDye 800CW Donkey anti Rabbit | LI-COR Biosciences | Cat# 926-32213, RRID:AB_621848 |
| **Chemicals, peptides and recombinant proteins** | | |
| Streptavidin | Sigma | Cat# S4762 |
| KO-DMEM | Gibco | Cat# 10829-018 |
| FBS | Pan Biotech | N/A |
| NEAA | Gibco | 11140-035 |
| L-glutamine | Gibco | 25030024 |
| Peniciliin/streptomycin | Gibco | 15070063 |
| Sodium Pyruvate | Invitrogen | 11360070 |
| 2-mercaptoethanol | Gibco | 31350010 |
| LIF | EMBL Protein Expression and Purification core facility | N/A |
| PD0325901 | Cambridge Bioscience | SM26-10 |
| CHIR99021 | Cambridge Bioscience | SM13- 50 |

| Reagent or resource | Source | Identifier |
|---|---|---|
| 0.5% Trypsin | Life Technologies | 25300054 |
| DMEM | Life Technologies | 11995065 |
| 4-hydroxytamoxifen | Sigma | H7904 |
| R0 | Sigma-Aldrich | A8094 |
| K0 | Sigma-Aldrich | L8662 |
| R10 | CK Isotopes | CNLM-539-H1 |
| K8 | CK Isotopes | CNLM-291-H |
| Dialysed FBS | Gemini Biosciences | FSD500 |
| DMEM for SILAC | Life Technologies | A33822 |
| OptiMEM | Gibco | 309600 |
| Alexa Flour 647 azide | Invitrogen | A10277 |
| Alexa Flour 488 azide | Thermo Scientific | A10266 |
| a-P32-dATP | Perkin Elmer | BLU012Z |
| EdU | Invitrogen | A10044 |
| BrdU | Sigma | *B5002* |
| Rapamycin | Cambridge Bioscience | SM83-5 |
| 5-ph-IAA | Cambridge Bioscience | HY-134653 |
| Nocodazole | Sigma | M1404 |
| Thymidine | Sigma-Aldrich | T9250 |
| RNase A | Sigma | R5250 |
| 20XSSC | life technology | AM9765 |
| Formamide | Sigma | 47671 |
| Formamide | Sigma | F9037A |
| Cot1 | Life technology | 18440016 |
| Salmon Sperm DNA | Invitrogen | AM9680 |
| EDTA-free protease inhibitor | Sigma | 5056489001 |
| Phosphatase Inhibitor Cocktail 3 | Merck Life Sciences UK | P0044-5ML |
| Biotin-azide | Thermo Fisher | B10184 |
| Puromycin | Sigma Aldrich | P-8833 |

| Reagent or resource | Source | Identifier |
|---|---|---|
| Blasticidin | Invitrogen | R210-01 |
| Paraformaldehyde | Sigma-Aldrich | P6148 |
| Vectorshield | Vector Laboratories | H-1900 |
| Vectorshield with DAPI | Vector Laboratories | H-1200 |
| Formaldehyde | Sigma-Aldrich | 252549 |
| Protein G dynabeads | Invitrogen | 10004D |
| proteinase K | Sigma-Aldrich | P6556 |
| H3 peptide | EpiCypher | cat # 12-0001 |
| H3K9me3 peptide | EpiCypher | cat # 12-0012 |
| H3K9me3S10ph peptide | EpiCypher | cat # 12-0054 |
| H3K9me1S10ph peptide | EpiCypher | cat # 12-0093 |
| H3K9me2S10ph peptide | EpiCypher | cat # 120155 |
| H3S10ph peptide | EpiCypher | cat # 12-0041 |
| Streptavidin beads | Thermo Fisher | 65002 |
| **Critical commercial assays** | | |
| Lipofectamine 2000 | Thermo Scientific | 11668019 |
| Click iT Cell Reaction Buffer Kit | Thermo Scientific | C10269 |
| **NEBNext® Ultra™ II DNA Library Prep Kit** | NEB | E7103L |
| NEBNext® Ultra™ II DNA Library Prep Kit | NEB | E6240 |
| *DIG NICK TRANS MIX* | Roche | 11745816910 |
| *Biotin NICK TRANS MIX* | Roche | 11745808910 |
| *Random primed DNA labelling kit* | Roche | 11004760001 |
| QuantaRed™ Enhanced Chemifluorescent HRP Substrate Kit | Thermo Scientific | 10053933 |
| TMT10plex TM Isobaric Mass Tag Labeling Kit | Thermo Fisher Scientific | 90113 |

| Reagent or resource | Source | Identifier |
|---|---|---|
| ChIP DNA clean and concentrator kit | Zymo research | D5205 |
| Qubit dsDNA HS assay kit | Invitrogen | Q32854 |
| Qubit dsDNA BR assay kit | Invitrogen | Q32850 |
| SPRIselect® Reagent Kit | Beckman Coulter, Inc. | B23317 |
| LightCycler® 480 SYBR Green I Master | Roche | 4887352001 |
| EZQ™ Protein Quantitation Kit | Invitrogen | R33200 |
| **Deposited data** | | |
| H3K9me3 ChIP-seq | This study | GEO: GSE299608 |
| Hi-C | Gnan et al., 2021 | GEO: GSE148244 |
| Repli-seq (*Rif1* wildtype and knockout cells) | Gnan et al., 2021 | GEO: GSE148244 |
| Repli-seq (O6 region tethering) | This study | |
| Suv39H1/2 HiC | Fukuda et al., 2021 | GEO: GSE169106 |
| H3K9me3 ChIP-seq Suv39h1/2double null | Aydan Bulut-Karslioglu et al., 2014 | GEO: GSE57092 |
| MS data | This study | |
| SILAC-MS | This study | |
| **Experimental models: cell lines** | | |
| *Rif1*$^{wt/wt}$: B, E, F, H | Foti et al., 2016 | N/A |
| *Rif1*$^{flox/flox}$*:* 5, 18, 20, 24, 28 | Foti et al., 2016 | N/A |
| *Rif1*$^{TgWT/flox}$: A7, G1, H4, H6 | Gnan et al., 2021 | N/A |
| *Rif1*$^{DPP1/flox}$: F8, G11, H1, H2 | Gnan et al., 2021 | N/A |
| *Rif1*$^{wt/flox}$*:* 19, 22 | Unpublished | N/A |
| **Rif1-mAID-11** | This study | N/A |
| **O6-Tether** | This study | N/A |

| Reagent or resource | Source | Identifier |
|---|---|---|
| **Recombinant DNA** | | |
| px458 | Addgene | RRID:Addgene_48138 |
| pMK381 | Addgene | RRID:Addgene_140536 |
| pMK346 | Addgene | RRID:Addgene_121180 |
| SP202 | Gift from Steven Pollard | N/A |
| SP208 | Gift from Steven Pollard | N/A |
| pACMJYP-Lap2B-WT-NL | Gift from Wendy Bickmore | N/A |
| pLyn11-FRB-mCherry | Gift from Susan Rosser | N/A |
| pSB1195 | This study | N/A |
| pSB1196 | This study | N/A |
| pSB1204 | This study | N/A |
| pSB1210 | This study | N/A |
| pSB1395 | This study | N/A |
| pSB1401 | This study | N/A |
| pSB1432 | This study | N/A |
| pSB1476 | This study | N/A |
| pSB1477 | This study | N/A |
| pSB1485 | This study | N/A |
| pSB1486 | This study | N/A |
| BAC: RP23-410A9 | BACPAC Resources | *N/A* |
| BAC: RP23-414K22 | BACPAC Resources | *N/A* |
| BAC: RP24-318O6 | BACPAC Resources | *N/A* |
| BAC: RP24-127L5 | BACPAC Resources | *N/A* |
| BAC: RP23-294E8 | BACPAC Resources | *N/A* |
| BAC: RP23-325L9 | BACPAC Resources | *N/A* |
| **Software algorithms** | | |
| Flowjo | V10.10.0 | FlowJo (RRID:SCR_008520) |
| RStudio | 2024.09.0+375 | RStudio (RRID:SCR_000432) |

| Reagent or resource | Source | Identifier |
|---|---|---|
| Jupyter notebook | V7.0.8 | https://jupyter.org/ |
| Integrative Genomics Viewer | Robinson et al., 2011; Thorvaldsdottir et al., 2013 | RRID: SCR_011793 |
| Galaxy | 25.0.2.dev0 | RRID:SCR_006281 |
| GraphPad Prism | 10.5.0 | RRID:SCR_002798 |
| ImageJ | | RRID:SCR_003070 |
| FastQC | V0.11.9 | RRID:SCR_014583 |
| Bowtie | | RRID:SCR_005476 |
| Samtools | V1.17 | RRID:SCR_002105 |
| DeepTools | | RRID:SCR_016366 |
| BEDTools | | RRID:SCR_006646 |
| MaxQuant | | RRID:SCR_014485 |
| Perseus | | RRID:SCR_015753 |
| Snakemake | V6.15.1 | RRID:SCR_003475 |
| Cutadapt | V1.18 | RRID:SCR_011841 |
| MultiQC | V1.26 | RRID:SCR_014982 |
| BWA | V0.7.16 | RRID:SCR_010910 |
| Picard MarkDuplicates | V3.3.0 | RRID:SCR_006525 |
| DiffBind | 3.8.4 | RRID:SCR_012918 |
| RegioneR package | 1.3.0 | https://bioconductor.org/packages/regioneR/ |

Table 2: Oligos

| Genotyping | Forward (5’-3’) | Reverse (5’-3’) | Notes |
|---|---|---|---|
| O6_scrn | ATTAGCCTGGGAAGGAGCTC | AACACAATCCCGACCTCCAT | To screen clones with attP1-attP2 landing pad integration to O6 loci. Non-targeted 197bp; targeted: 275bp |
| TetO_scrn | ATTAGCCTGGGAAGGAGCTC | CGTGATACGGATCTCGAGG | To screen clones with TetO arrays integrated. TetO integrated: 180bp |
| Rif1 | TTAGAGGAACTGAGGGGTAGGTAG | AACTGCAACTCTACTGAGGGAAG | To genotype cells after OHT treatment. Rif1 wtt: 78bp; flox: 138bp; deleted: 246bp |
| | | TGAAACCGTAGCCAGAAACTG | |
| RFHF-WT | TCTGTTAGCAACGCGACCTT | TTTACGAGCTTTAGGCGGCA | To genotype Rif1 tgWT, 200bp |
| RFHF-DPP1 | AGATTCGCCGTGTCAGCGCC | TGATCGGGCTGCTATTGCTGG | To genotype Rif1 DPP1, 110bp |
| Rosa26 scrn | GCCAGTCCAAGAGAAAGCAC | AGAATCCCTTCCCCCTCTTC | To screen clones with Rosa26 alleles targeted. WT: 1144bp; targeted: 1505bp |
| | | AGCTTGGCGTAATCATGGTC | |
| AID-Rif1 scrn | GGTTACTGGGTGTCTTCCGG | GTAATCTCAACCACTCCGGCAG | To screen clones with Rif1 targeted. WT:533bp; targeted: 215bp |
| | | TGCTCCGTCCATTGATACCTTC | |
| **Southern probe** | **Forward (5’-3’)** | **Reverse (5’-3’)** | **Notes** |
| O6 _Southern | TTTGCTGTCCCTACGATTACG | CTTCCCACTATAGCACTGAGCT | To PCR genomic fragment to use as probe. 514bp |
| RIF1FLAGHA_southern | GTGTCACTACTCTCACATTT | TGTTTTTCCATTTAGAAGCCAG | To PCR southern blot probe R5. 848bp |
| **CRISPR-guide** | **Top** | **Bottom** | **Target site** |
| Rif1_sg1 | CACCGCGGCCGACATGACGGCCCC | AAACGGGGCCGTCATGTCGGCCGC | GCGGCCGACATGACGGCCCCagg |
| Rif1_sg2 | CACCGCGACCTGGGGCCGTCATGT | AACACATGACGGCCCCAGGTCGC | GCGACCTGGGGCCGTCATGTcgg |
| O6 | CACCGCAACACAATGCGCGCACAT | AAACCATGTGCGCGCATTGTGTTGC | GCAACACAATGCGCGCACATcgg |
| Rosa26 targeting | CACCGGCCCATCTTCTAGAAAGAC | AAACGTCTTTCTAGAAGATGGGC | GCCCATCTTCTAGAAAGACtgg |
| **Gblocks** | | | |
| Rif1 5’ homology | gtaaataagcgcgagccgggagcggacggcgggccccggggcggcgagctgcggagcggacttcccgcgtggggggtgag [illegible] | | |
| Rif1 3’ homology | gtgaggcgggccggggggacgagcccggggcgggagccggggcagccgtggggacccggcggacgcgggtagttcgggg [illegible] | | |
| **ChIP-qPCR primers** | **Forward (5’-3’)** | **Reverse (5’-3’)** | **Notes** |
| O6 | GCTAGCCACCACAGGAAAAG | GCTCAGACACCCTATCCCTC | 86bp; 60˚C |
| TetO site | ATAAGCTTGATGCTAGCCGG | GAAACACAATCCCGACCTC | 114bp; 60˚C |
| B-actin | ATGAAGAGTTTTGGCGATGG | GAGACATTGAATGGGGCAGT | 91bp; 60˚C |
